# Response of soil and vegetation in a warm-temperate Pine forest to intensive biomass harvests, phosphorus fertilisation, and wood ash application

**DOI:** 10.1101/2021.03.16.435291

**Authors:** Laurent Augusto, Florent Beaumont, Christophe Nguyen, Jean-Yves Fraysse, Pierre Trichet, Céline Meredieu, David Vidal, Valérie Sappin-Didier

## Abstract

**Background and Aims:** Concerns about climate change and carbon economy have prompted the promotion of alternative energy sources, including forest-based bioenergy. An evaluation of the environmental consequences of intensive harvests (stumps and roots, and also branches and foliage) for energy wood supply, and use of wood-ash recycling as a compensatory practice, helps in the evaluation of the use of forest biomass for energy production.

**Methods:** We made use of records from a split-plot experimental site crossing four different intensities of biomass harvesting (Stem-Only Harvest [SOH], Aboveground Additional Harvest [AAH], Belowground Additional Harvest [BAH], and Whole-Tree Harvest [WTH]) and three compensation methods (control [C], wood ash application [A] and phosphorus fertilisation [P]) to evaluate, in the 11-years-old stand (maritime pine: *Pinus pinaster*) that followed the biomass exports of the former stand, their effects on nutrient budgets, tree growth, soil fertility, chemical properties and soil carbon. This site is located in a forest on a poor soil, under a warm temperate climate (SW France).

**Key results:** Despite their low additional biomass exports (+10% for AAH to +34% for WTH), the non-conventional harvest practices exported much higher quantities of nutrients than the conventional SOH technique (for example +145% for N and K in WTH). Consequently, these treatments had negative effects on the soil nutritive status. Additional biomass harvests impacted the soil organic matter content, with negative effects on P_-organic_, soil cation exchange capacity, exchangeable Ca, and most extractible nutrients. However, data suggested that tree growth and foliage nutrient content had not yet been significantly impacted by harvest treatments, whereas tree nutritional status was improved by P-fertiliser or wood ash. As expected, we observed a positive effect of wood ash application on soil pH and nutrient content but, like additional harvests, wood ash application decreased the pool of soil organic carbon (~10% of the initial stock with ~7% of N_-total_ losses).

**Conclusions:** Overall, this factorial experiment showed that exporting more forest biomass due to the additional harvesting of tree canopies, stumps and roots had negative consequences on the ecosystem biogeochemistry. Additional harvests have aggravated the poverty of the already oligotrophic soil, and decreased the soil organic carbon content. Importantly, applying nutrients as fertiliser or wood ash did not compensate for all the negative impacts of biomass exports and the method of wood ash recycling in forests could even decrease the soil organic carbon.

## Introduction

Although there is a consensus that forests constitute an important leverage for climate change mitigation (IPCC 2014), there is also a heated debate on how forests should be managed for mitigation purposes (Lindner and Karjalainen 2007). In particular, it is debated to what extent additional harvesting of forest biomass, such as foliage, branches, stumps, and roots (Nicholls et al. 2009) should be promoted for energy production. Indeed, this practice can impoverish forest ecosystems because these tree compartments dedicated to energy wood supply −the so-called *harvest residues* or *logging residues*– are rich in nutrients (Andre et al. 2010; Achat et al. 2015a; Augusto et al. 2015a). Review studies indicate that intensive biomass harvesting −as compared with conventional harvesting− can have negative consequences on ecosystem functioning, such as soil nutrients and organic matter pools, and consequently on future forest growth (Thiffault et al. 2011; Wall 2012; Achat et al. 2015a). Extracting more biomass from forests without any compensatory fertilisation seems to be an unsustainable leverage for climate change mitigation (Garcia et al. 2018). Therefore, it has been proposed that intensive harvests should be associated with compensatory practices (Nohrstedt 2001; Ranius et al. 2018; Ventura et al. 2019).

Applying wood ash is often promoted as a good method to compensate for the negative effects of intensive harvests on the ecosystem nutrient budget (Hannam et al. 2018; Ranius et al. 2018; Ventura et al. 2019). Indeed, wood ash has a high nutrient content (Aronsson and Ekelund 2004), can reduce soil acidity (Reid and Watmough 2014), and contributes to the concept of *circular bioeconomy* as wood ash returns some of the nutrients that were exported by the removal of logging residues (Pitman 2006). On the other hand, applying wood ash might contaminate ecosystems because of its relatively high content in several micronutrients or non-essential metals (Vance 1996; Nnadi et al. 2019). Unfortunately, while several meta-analyses have been carried out on intensive biomass harvests (Thiffault et al. 2011; Achat et al. 2015a) or wood ash application (Augusto et al. 2008a; Reid and Watmough 2014), a combination of these two practices have rarely been studied together to assess their interactive consequences (Hagerberg and Wallander 2002; Wang et al. 2010). Therefore, we conducted a field experiment whose main objective was to evaluate these possible interactions.

Located in south-western France, the *Landes de Gascogne* forest (where our study took place) has several advantages for evaluating the environmental consequences of intensive harvests and their compensatory practices. Firstly, it is a large man-made pine forest almost entirely dedicated to intensive forestry. As such, the area is already subjected to collection of harvest residues (mainly stumps and roots, and to a lesser extent also branches and needles (Mora et al. 2014; Banos and Dehez 2017)) for energy wood consumption (Augusto et al. 2010, 2015a), which in turn produces large amounts of wood ash (Alvarez-Alvarez et al. 2018). Secondly because this forest is strongly oligotrophic (Augusto et al. 2010), and particularly poor in phosphorus (Achat et al. 2009), it has been diagnosed to be particularly sensitive to the potential negative consequences of intensive harvest (Durante et al. 2019). Finally, it is a warm–temperate forest. Indeed, knowledge about wood ash application and removal of logging residues is mainly based on studies carried out in boreal forests (Augusto et al. 2008a; Walmsley and Godbold 2010), whereas the consequences of these management practices are probably climate-dependent (Achat et al. 2015b). For instance, several Nordic studies indicated that stump harvests, or wood ash application, had no effect on soil organic carbon (SOC) while studies conducted in warmer latitudes concluded that the impact was highly negative (Zabowski et al. 2008; Stromgren et al. 2013; Jurevics et al. 2016).

Our initial expectations were based on the current knowledge about intensive biomass harvest or wood ash application, in comparison with the local conditions of climate, soils, and forestry. More explicitly, we expected the following effects (Table 1).

**Table 1.** Initial expected consequences of intensive biomass harvesting and wood ash application on the studied ecosystem (temperate oligotrophic forest) (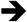: means “has no effect on…”; 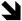: means “decreases…”; 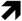: means “increases…”; **?**: means “unknown effect on…”)

| Ecosystem trait |  | Intensive harvests | Wood ash |
| --- | --- | --- | --- |
| Soil fertility<br>(phosphorus)<br>{hypothesis H1} | *General knowledge* | ⬇ soil P | ➔ soil P |
|  | *Local conditions* | P-limited soils |  |
|  | *Expected results* | ⬇ (aggravated P-limitation) | ➔ (or ⬆) soil P |
| Tree growth<br>(carbon sequestration)<br>{H2} | *General knowledge* | ⬇ tree growth | ➔ (or ⬆) tree growth |
|  | *Local conditions* | oligotrophic P-limited ecosystem |  |
|  | *Expected results* | ⬇ tree growth | ➔ tree growth |
| Soil Organic Matter<br>SOM<br>(carbon and nitrogen)<br>{H3} | *General knowledge* | ➔ boreal forests<br>⬇ temperate forests | ➔ boreal forests<br>? temperate forests |
|  | *Local conditions* | warm temperate |  |
|  | *Expected results* | ⬇ SOM | ➔ SOM |
| Soil acidity<br>(pH, base saturation)<br>{H4} | *General knowledge* | ⬇ acido-basic status | ⬆ acido-basic status |
|  | *Local conditions* | acidic soils |  |
|  | *Expected results* | ⬇ acido-basic status | ⬆ acido-basic status |
| Trace metals in<br>ecosystem<br>(micronutrients:<br>[B, Cu, Fe, Mn, Mo, Zn]<br>and toxic trace metals<br>[As, Pb, Cd])<br>{H5} | *General knowledge* | ➔ or ⬇ trace metals<br>(due to exports) | ➔ for podzol soils<br>(boreal soils in Nordic areas) |
|  | *Local conditions* | low content in tree biomass growing on podzol soils |  |
|  | *Expected results* | ➔ trace metals | ➔ or ⬇ bioavailability<br>at moderate ash dose<br>+ increase pH |

Hypothesis H1: the soil phosphorus availability, the most limiting nutrient in the sandy acidic soils of the study region, would be decreased by intensive harvests (Achat et al. 2015a) and would be only partly compensated by wood ash that has a low phosphorus content and low bioavailability (Steenari and Lindqvist 1997; Fransson et al. 1999).

H2: the sequestration rate of C into forest biomass (estimated as the growth rate of the subsequent forest stand) would be decreased by intensive harvests (Achat et al. 2009) because the local soils are nutrient poor (Achat et al. 2015b).

H3: the size of the soil organic carbon content would be decreased by intensive harvests as for most temperate forests (Achat et al. 2015a, b), but might remain unaffected by wood ash application as observed in boreal forests (Augusto et al. 2008a).

H4: the soil would be acidified by intensive harvest, but partly neutralised by wood ash (Reid and Watmough 2014; Achat et al. 2015a).

H5: because the local pine trees contain small amounts of trace metals in their biomass compared to stocks in the soil (Saur et al. 1992; Trichet et al. 2018), exports through biomass harvest and returns through wood ash application would not significantly change the trace metal content in the ecosystem.

## Materials and Methods

### Study region and experimental site

The *Landes de Gascogne* area is a flat region ~1.3 Mha in south-western France (between 43.5 and 45.5°N, and 1.5°W and 0.3°E). The climate is warm oceanic (mean annual temperature = 13°C, with mean monthly values ranging from 6.0 to 20.3°C), usually with a dry season in summer and a wet season in winter and/or in spring (annual rainfall is around 900 mm per year). Soils are sandy podzols (9–14% fine sands; 80–89% coarse sands), which are classified as more or less hydromorphic and humic (*entic podzol* in the WRB soil classification), or with a hardpan (*ortsteinic podzol*; Augusto et al. (2010)). They developed in Aeolian deposits of quaternary sands (the *Sables des Landes*) and are characterised by their coarse particle size and their high quartz content (Bertran et al. 2011). Soils are acidic and poor in nutrients, particularly in phosphorus (P) and they have been identified as being among the poorest soils in P in the world (Achat et al. 2009; Bredoire et al. 2016). The presence of wetlands is the consequence of the low drainage capacity of the landscape (low elevation plain with streams but only one river) that makes the watertable close to the soil surface (Deirmendjian et al. 2018). Initially grown for its resin, Maritime pine (*Pinus pinaster* Ait.) is now used for wood production and is often P-fertilised at plantation to alleviate the main nutritional limitation of the region (Trichet et al. 2009), and stands are surrounded by ditches to improve water drainage in rainy periods.

### Experimental design and field procedures

The area used for this study is called the *Forêt du Nezer* (44.57°N, 1.05°W; elevation = 21 m asl; slope < 1%; entic podzol). The experimental site is a split-plot factorial design, crossing four different intensities of *Biomass Harvesting* (BH factor in main plots) and four *Compensation Methods* by applying wood ash or phosphorus fertilisation (CM factor in subplots nested in BH plots), resulting in sixteen treatments replicated in three blocks (Figure S1). Contrary to the CM treatments, the BH treatments had to be applied to large plots (main plots) because they required the use of large machines. In April 2007, the standing forest (45-year-old, 465 trees ha^-1^; basal area = 29.7 m^2^ ha^-1^) was clear-cut. In practice, twelve large square plots, 100 × 100 m (1 ha large), were established in order to apply the three blocks × four *Biomass Harvesting* treatments, described below:

1. Stem-Only Harvest (‘SOH’; stem harvest down to 7 cm of diameter above bark) without any residue harvest (exports = stems: this has been the local conventional practice for decades),
2. Aboveground Additional Harvest (‘AAH’), which included stem harvest with additional harvesting of branches with part of their foliage (see below; exports = stems + canopies; canopy = branches + twigs + foliage),
3. Belowground Additional Harvest (‘BAH’), which included stem (without branches, twigs, and foliage) harvest with additional harvesting of stumps and roots (exports = stems + stumps + roots),
4. Whole-Tree Harvest (‘WTH’) with additional harvest of all other tree compartments (exports = stems + branches + twigs + foliage + stumps + roots).

The conventional harvesting (clear-cut) was carried out in April 2007 (see Figure S2 for more details about the study sequence), with particular attention to the slash (branches and needles) which was gathered in each of the four 1 ha plots (Figure S3a). In June 2007, the branches were bundled, forwarded from the six main plots (AAH-WTH plots × 3 blocks; Figure S3b-d), and finally transported to the power plant. The time-lapse between this harvest and the clear-cut has enabled a partial fall of the foliage (needles, in unquantified proportions). Then, in July-August 2007, the stumps were excavated with their main roots, fragmented into a few pieces, and left on the ground in the six main plots (BAH-WTH plots × 3 blocks; Figure S3e-g), enabling the natural removal of the soil particles by the rain. These fragments were harvested in February 2008. Two years after clear-cutting, in January 2009, the area was strip ploughed to a depth of 0.3 m. The ploughed strips were repeated every 4 meters and dedicated to seedling plantation. The soil surface of the ploughed strips (representing ~half of the total surface area) was flatten with discs before the plantation of 1,250 seedlings per hectare (4 × 2 m) of one-year-old maritime pine seedlings of local origin.

Forty-eight small sub-plots, 50 × 50 m (0.25 ha), were established within the two-year-old plantation. Four sub-plots of compensation methods (CM factor) were nested in every main plot (dedicated to a BH treatment; Figure S1). The *Compensation Methods* were applied in July 2011 for the wood ash and in December 2011 for the other fertilisers, as follows:

1. Control: no nutrient application (‘C’),
2. Phosphorus fertilisation (‘P’), considered as standard in this forest area (application of 60 kg-P_2_O_5_ per hectare in the form of superphosphate (30% P_2_O_5_)),
3. Phosphate-potassium fertilisation (the same P treatment as above with the addition of 60 kg-K_2_O per hectare in the form of potassium sulphate (50% K_2_O)). It should be noted that this CM treatment was not included in this study.
4. Wood ash application (‘A’) consisted of the application of 5 Mg of loose ash per hectare (provided by a local power biomass plant). The ash composition is shown in the Table S1.

### Plant measurements and field sampling

#### Estimates of biomass and nutrient exports (*Biomass Harvesting* factor)

The quantity of biomass, and associated nutrients, exported from the plots due the Biomass Harvest treatments were estimated, based on a combination of measurements and models. In practice, before clear-cutting the standing pine stand, all trees were identified and each diameter was measured at breast height (≈ 1.3 m).

##### Biomass estimates

Then, we selected 14 trees that represented the range of stem size of the forest. All these trees were cut down and studied intensively:

i. the stem diameter was measured, and wood disks were sampled, at several heights (0 m, 0.5 m, 1.3 m, and then every 2 m); the wet mass and the dry mass of the stem disks (wood and bark, separately) were measured,
ii. the diameter of all branches was measured at 10 cm from their insertion level,
iii. the total dry mass of three branch classes (living branches, treetop [*i. e*. the upper part of the stem, at diameter < 7 cm], dead branches) were weighed (all branches of each class were pooled),
iv. the dry mass of 5 living branches (wood + bark + foliage) per tree was weighed, and
v. the dry mass values of the stem wood, and the stem bark, were measured separately.

The total dry mass by compartment of each sampled tree was then calculated. The tree size measurements (diameter at breast height) were used to estimate the aboveground biomass of the different tree compartments using allometric relationships (unpublished FCBA models, calibrated with 45 trees distributed in 5 other pine stands of a similar age). The estimated biomass values of the tree compartments were consistent with the measured values, so the models were used to calculate the biomass at the plot scale, based on the inventory of all trees.

The <u>effective export rate</u> of the different tree compartments was estimated in different ways, depending on the compartment studied:

i. stem wood was assumed to be harvested in totality (export rate = 100%),
ii. stem bark harvests were quantified by collecting all the bark that remained on the floor of a given area, after the stems exported from five other stands of the region (FCBA unpublished data; export rate ≈ 80%),
iii. the effective exports of the canopies (*i.e*. branches + twigs + foliage) and of the tree belowground (stump + roots) were estimated by direct measurements of the harvests realised. In practice, the harvested canopies were gathered into ~2.5 m long bundles (n = 465 in total), labelled per plot, and taken to the local power biomass plant. Bundles were weighed individually (fresh weight = 225-253 kg; mean weight = 240 kg), and 20 bundles were taken at random to estimate their dry matter content (mean dry content = 67% of the initial mass). All bundles were crushed to provide pellets for the power plant. During the crushing stage, 36 samples of biomass were taken at regular time intervals to estimate the dry matter content. This procedure enabled us to estimate the effective harvest rate of the canopy.

Conversely, it was not logistically feasible to measure the effective rates for foliage, twigs, and large branches. Based on visual observations and on field studies (Stupak et al. 2008), we assumed that (i) the twig harvest rate was 30% higher than that of the foliage, and (ii) the branch harvest rate was 50% higher than that of the twigs. Finally, based on these ratios and on the measured harvest rate of the whole canopy, the harvest rate values were adjusted. The values of the nutrient content in the different canopy compartments (*i.e*. needles, twigs, branches, stembark, stemwood) were the mean values measured on samples from the 45 trees used to build allometric relationships (see above). These mean values were within the ranges commonly observed in the study region (Augusto et al. 2008b; Trichet et al. 2018).

A similar approach to that used for branches was used for the belowground biomass. In practice, the potentially available belowground biomass (*i.e*. stump + roots) was estimated for each tree based on allometric relationships that use the stem diameter at breast height as predictive variable (Augusto et al. 2015a). The realised belowground biomass harvest was estimated from the weight of all the fresh biomass per plot (by weighing the trucks, with and without the transported biomass). The water content of the biomass was measured on 36 samples in order to calculate the dry biomass (Figure S3g; sampling at regular intervals during the crushing flow; mean value per plot of the dry content = 65-72%). These samples were used also for nutrient analyses. The measured values of nutrient content in stumps and roots were within the ranges commonly observed (Augusto et al. 2015a), except for the potassium content that was higher in the present study (1.50 versus 0.99 mg g^-1^).

All these measurements and analyses enabled us to quantify the standing biomass before the Biomass Harvest treatments, and the effective exports due to harvests in terms of biomass and nutrients.

### Field measurements in the subsequent forest plantation

#### Tree growth and needle sampling

The growth rate of the new plantation −established after biomass harvest− was assessed simply by measuring the standing biomass of the young trees. Indeed, because the survival rate of the planted seedlings was nearly 100%, and not affected by experimental treatments, the stand density was the almost the same in all subplots. The individual size of trees was estimated by measuring 50 pines, chosen at random, per subplot. Tree height and stem circumference at breast height were measured in December 2015 and 2019 (*i.e*. on 7- and 11-year-old trees). The aboveground biomass was then estimated based on allometric relationships (Vidal et al. 2019).

Needles were sampled to assess the nutritive status of trees, and their degree of contamination by trace elements. For this, 8 pines were first selected at random in each subplot. Current-year needles were collected in December 2019 using a telescopic pruner. Needles were collected in the upper third of the canopy, where sunlight is always directly available, and in two opposite directions. The needles were cleaned in a bath of demineralised water, pooled by subplot, and then dried in an oven for ten days at a temperature of 40°C.

#### Understorey survey

The understorey composition was assessed in May 2019, using the *“phytovolume”* approach (Gonzalez et al. 2013; Vidal et al. 2019). In practice, the same pair of operators surveyed all the surface area of the studied subplot, and then estimated the soil cover percentage of the main plant functional types (*i. e*. the perennial herb *Molinia caerulea*, the bracken fern *Pteridium aquilinum*, ericaceous shrubs (*Calluna vulgaris* and *Erica scoparia*), and the common gorse *Ulex europaeus*). The mean height of each plant functional type was estimated based on three height measurements representative of the plant height range in the surveyed area (Figure S4). The phytovolume value was computed as the product of the ground cover (estimated in squares of 16 m^2^) and the plant height. Finally, the phytovolume value was converted into estimates of plant aboveground biomass using dedicated allometric relationships that were previously calibrated with destructive measurements of the standing biomass (Gonzalez et al. 2013; Vidal et al. 2019).

#### Soil sampling

In December 2019, a composite sample of the topsoil was built in each of the 36 subplots. To do this, a systematic grid of 8 sampling points was used per subplot: 4 points on the tree ridges, and 4 points in the furrows, to capture the spatial distribution of the topsoil microtopography induced by the regular design of the strip ploughing (Forrester et al. 2013) and tree plantation and by subsequent operations of understorey control (*i.e*. a bladed roller passing in the furrows). Because of the soil preparation before tree plantation and because stands were still young, the forest floor layer was thin at soil sampling. We consequently did not sample it, although we recognize its importance for plant nutrition (Jonard et al. 2009). After having gently removed the forest floor layer, we sampled the topsoil (0-15 cm layer) using a corer (diameter = 8 cm). All samples were bulked in the field, and continuously homogenised by hand until the composite sample showed no heterogeneity. The soil bulk density (kg L^-1^) was estimated based on a pedo-transfer function specifically calibrated for the local soils (Augusto et al. 2010).

### Laboratory analyses

#### Plant analyses

The needle samples were ground (< 1μm) using a mill in titanium to avoid contamination of the sample. Aliquots of powdered samples (0.25g) were digested using a mixture of 1 ml HNO_3_ and 4 ml H_2_O_2_ in DigiPREP System in dry bath blocks. Then, the solution was filtered to 0.45 μm and the concentrations of K, Ca, Mg, Na, Fe, Mn, Al, Cr, Cu, Ni, Zn, and Co in the extracts were analysed by ICP-AES and Cd, Pb, Tl, and Mo by ICP-MS. The validity and accuracy of the procedures were checked using a standard reference material (SRM-1573a, NIST).

#### Soil characteristics

Soils were sieved to < 2 mm and air-dried. All analyses (except pH) were carried by INRAE soil testing laboratory (INRAE-LAS, Arras, France), according to French standardised procedures or international procedures. Soil pH was determined in a distilled water (1:5 ratio) and 0.01 M CaCl_2_ extract (1:10 ratio; standard NF ISO 10390:2005). The soil organic carbon content (SOC) and total nitrogen (N_-total_) were determined by dry combustion after correction for carbonate (SOC: NF ISO 10694:1995, N_-total_: NF ISO 13878). The available phosphorus (P_-Olsen_) was determined using the Olsen method (NF ISO 11263). The cation exchange capacity (CEC) and exchangeable cations (K^+^, Ca^2+^, Mg^2+^, Na^+^, Fe^2+^, Al^3+^, and Mn^2+^) were measured at soil pH using the cobaltihexamine chloride method (NF X 31–130:1999). The soil texture was determined (NF X 31–107:1983), but only on three soil samples (composite samples corresponding to the three blocks of the site).

Total soil mineral element contents (major and trace) were quantified after solubilisation by fluorhydric and perchloric acids (NF X 31–147:1996). After complete dissolution, the concentrations of elements were determined by ICP-AES and ICP-MS (see above).

#### Soil solution extraction and analyses

Two chemical extractions were realised for the soils to estimate the availability of trace metals: ultra-pure water and ammonium nitrate 1M (NH_4_-NO_3_). Water extractions make it possible to estimate the content of elements that would be found in the soil solution. Ammonium nitrate 1 M extractions quantify the elements weakly adsorbed onto the solid phase by ion exchange at pH=6.

Ultra-pure water extraction was performed at the 1:5 soil to solution ratio, stirring on a roller-shaking table for one hour. At the end of the shaking, the solution was filtered (0.22 μm) and then part of the solution was acidified at 2% with HNO_3_ for trace metal analysis. The second part was reserved at 4°C to measure: pH, the concentration of dissolved organic (DOC) and inorganic (IC) carbon (by oxidative combustion; TOC-VCSH, Shimadzu), and anions (PO_4_^3-^, NO_2_^-^, NO_3_^-^) and NH_4_^+^ (colorimetric method; Technicon auto analyser II).

The NH_4_-NO_3_ extractions were performed at 1:2.5 soil to solution ratio (Symeonides and McRae 1977). The suspensions were shaken, filtered at 0.22 μm, and then acidified with 2 % HNO_3_. The concentrations of P, K, Ca, Mg, Na, Fe, Mn, and Zn in extracts from two extractions were assayed by ICP-AES and Cd and that of Pb and Cd by GF-AAS.

### Input-output budgets and data analyses

Estimating input-output budgets enables to evaluate if a forest management is sustainable (Ranger and Turpault 1999). However, because our study was not designed to quantify biogeochemical fluxes, the presented budgets should be considered with caution as they are only rough estimates, and further research is needed to assess the consequences of intensive forestry on ecosystem biogeochemistry. In practice, atmospheric deposition was estimated based on data from a nearby monitoring site (Croise et al. 2005), and values were adjusted to take into account the tree height effect on deposition (de Schrijver et al. 2012). Symbiotic nitrogen fixation at 11 years-old was estimated based on gorse biomass (see above), and taking into account the nitrogen content and fixation rate of this species (Cavard et al. 2007; Augusto et al. 2009). For periods after 11 years-old, which corresponds to the canopy closure of pine stands, a flux a 2.8 kg_-N_ ha^-1^ yr^-1^ was retained (Augusto et al. 2005). The flux of nutrients released by weathering of soil minerals was estimated based on soil mineralogy and the PROFILE model (Sverdrup et al. 2006). Elements that were added to the ecosystem through application of wood ash and fertiliser were quantified based on ash composition (Table S1) and fertiliser composition (unpublished data and Kabata-Pendias 2000). Exports of nutrients though biomass harvest were estimated by direct measurements (see above). Losses of nutrients through deep seepage were estimated based on an on-going monitoring experiment (XyloSylve) with the same tree species and similar tree age, soil, climate, and experimental treatments. Because input-output budgets should be calculated for a complete silvicultural rotation (Kimmins 1974), we calculated them not only for a stand age of 11 years-old (when we sampled plants and soil) but also at 20 and 40 years after plantation as it corresponds to the local rotation duration for intensive and conventional management, respectively. We did not present values of soil nutrient stocks, however it is possible to estimate soil pools (in kg_-element_ ha^-1^) by multiplying the soil content (in mg_-element_ g^-1^, which is equivalent to kg-element Mg^-1^) by 1578 (which is the mean value of soil mass of the 0-15 cm layer; in Mg_-soil_ ha^-1^; (Augusto et al. 2010)).

Data processing was performed with R, version 3.6.1. (R Core Team 2019). The split-plot design was analysed by a mixed effect model (*lme* function, package nlme 3.1-143). The biomass harvest factor and the compensation method factor were fixed effects (main and interaction effects). The block and biomass harvest factor nested in blocks were random effects associated with the intercept. When detected, heteroscedasticity was corrected by modelling the variance with the *varIdent*() function (nlme package) which allows different variances by level of the biomass harvest and compensation methods. Least-squares means were calculated (emmeans package 1.4.2) and linear contrasts were used to test for significant differences between treatments and their corresponding controls, namely SOH for the biomass harvest intensity (BH factor) and no fertilisation (C) for the compensation methods (CM factor). When necessary, the *P*=0.05 significance probability was adjusted for multiple pairwise comparisons (Tukey’s test).

Preliminary analyses of the results showed that the interactions between the BH factor and the CM factor were almost never statistically significant, as already been reported (Hagerberg and Wallander 2002), and that is why we analysed these factors separately.

Because of the lack of significant interactions between the two studied factors, the mean values of the Biomass Harvest treatments (n = 3 blocks = 3 replicates) pooled the values of the Compensation Methods subplots. Similarly, the mean values of the Compensation Methods treatments corresponded to 12 replicates (3 blocks × 4 BH treatments). The difference of number of replicates (3 versus 12), due to the split-plot design, implied that the statistical power is greater for the CM factor nested in BH compared to the BH factor. The main consequence of this split-plot design was consequently a greater difficulty in detecting significant effects of the BH factor.

Mean values are shown ± one standard error. The mean values of the main response variables are shown in Appendix 1, for the all 12 experimental combinations of treatments (4 BH × 3 CM). In addition, all data and scripts are available at https://doi.org/10.15454/LCU6OZ.

## Results

### Initial stand biomass and nutrient content, and effective harvests

The standing biomass of the stand prior to harvests was 179.1 ± 3.5 Mg ha^-1^ of which 40.1 ± 0.7 Mg ha^-1^ were belowground (Table S2). The standing biomass was distributed homogeneously among the experimental plots, with a coefficient of variation of 1% among the WTH plots to 8% among the SOH plots (CV=6% at the site scale). This biomass contained 24.7 ± 0.5 kg-P ha^-1^ (phosphorus) and 311 ± 6 kg-N ha^-1^ (nitrogen; Table S2). The harvesting treatments did not export all this biomass and its associated nutrients: the effective rate of harvest was measured as 71% for belowground biomass (in BAH and WTH treatments) and 91% for aboveground biomass (in AAH and WTH treatments). Within the aboveground biomass pool, the measured harvest rate of canopies (foliage + twigs + branches) was 53%, with estimated rates of 34%, 44%, and 66% for foliage, twigs and branches, respectively. The realised export of biomass was 115.32 Mg ha^-1^ for the conventional harvest (SOH), with moderately increased values for more intensive harvests (from +10% for AAH to +34% for WTH; Figure 1a).

**Figure 1.**
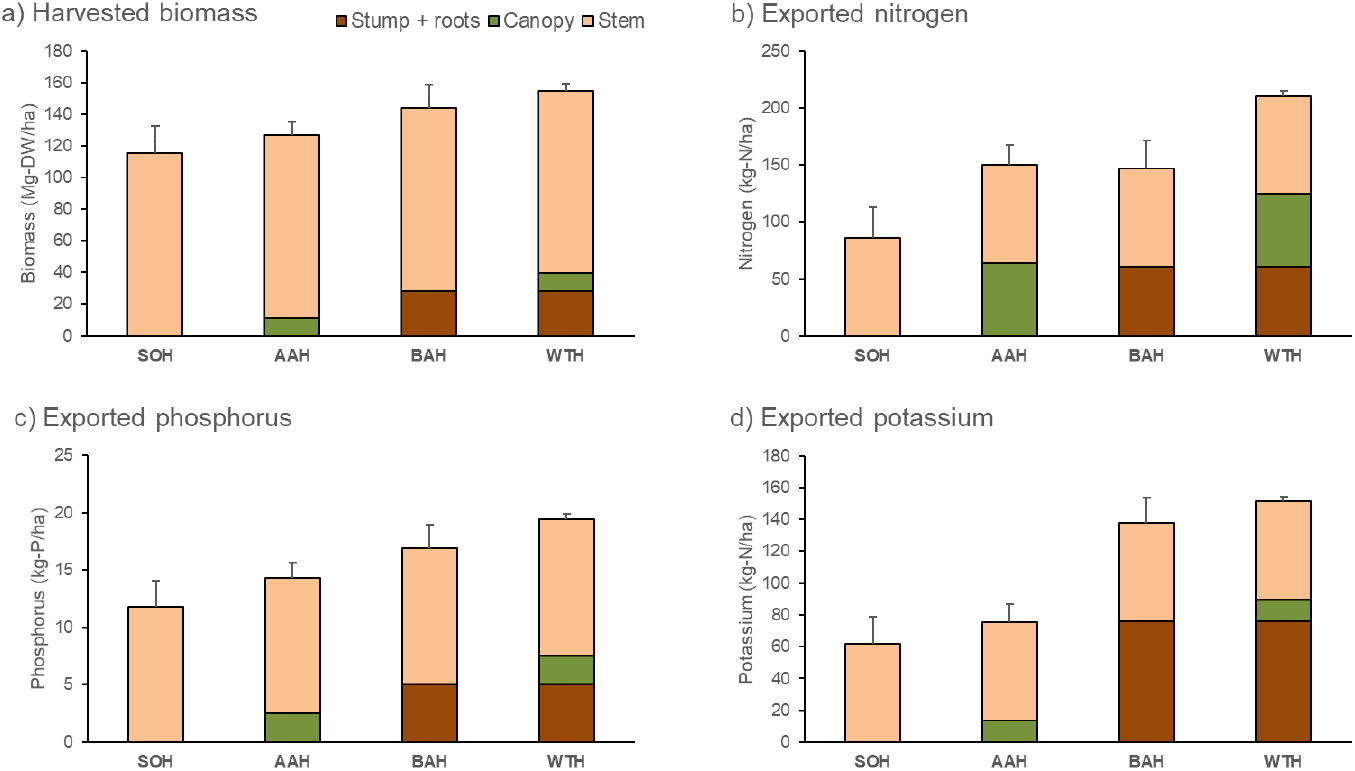
Realised exports of tree biomass and nutrients from the cut forest. SOH = stem-only harvest; AAH = aboveground additional harvest; BAH = belowground additional harvest; WTH = whole-tree harvest. Error bars are standard errors. All data (including data about calcium, and magnesium) are presented in Table S2. The methods used to quantify the biomass and nutrient exports are detailed in the manuscript in the section “*Plant measurements and field sampling*”, in particular in the subsection “*Estimates of biomass and nutrient exports*”.

Despite their low additional biomass exports, due to the differences in nutrient contents between tree parts, the non-conventional harvest practices exported much more nutrients than the conventional SOH technique. This was particularly the case for the canopy export of nitrogen (+74% in AAH; Figure 1b) and the belowground export of potassium (+123% in BAH). In the most impacting treatment (WTH), the increase in nutrient exports was between +64% for P (Figure 1c) and +145% for N and K (Figure 1b and 1d).

### Soil properties and fertility (hypothesis H1)

The main soil characteristics were representative of the sandy podzols of the region. For instance, the mean soil particle size distribution showed the dominance of sand (85.7±0.2%, 7.5±0.5%, 0.7±0.3%, 2.4±0.2% and 3.6±0.3%, for the coarse sand, fine sand, coarse silt, fine silt and clay fractions, respectively), and a low initial value of the total soil P content of 44±17 μg-P g^-1^ (see Augusto et al. (2010) for comparisons).

The biomass harvesting treatments had many effects on the soil nutritive status, but the effects depended on the form of the nutrients and micronutrients. While the AAH treatment showed increases in values of K, Na, Fe and Mn total contents (Table 2), the pattern was quite different for exchangeable cations (Table 3) and extractable nutrients (Table 4). Indeed, the nutrients that constitute the so-called *basic cations* of the CEC (*i. e*. K^+^, Ca^2+^, Mg^2+^, and Na^+^) were depleted to varying extents (except for K^+^) by the non-conventional harvest practices (AAH, BAH, and WTH) as compared with the SOH treatment (Table 3). The same pattern applied to Ca, Mg, and to a lesser extent K, that were NH_4_-NO_3_ extracted (Table 4). The losses in these nutrients (Ca, Mg, Mn, and Zn) were generally compensated by the application of wood ash (Table S1), as shown by the significant increases in their soil content as compared with the control treatment (Tables 3 and 4).

**Table 2.** Soil content in carbon, nutrients, and micronutrients as affected by biomass harvest and compensation methods

|  | SOC<br>(mg g<sup>-1</sup>) | DOC<br>(mg L<sup>-1</sup>) | N<br>(mg g<sup>-1</sup>) | C/N<br>unit less | P<sub>Olsen</sub><br>(μg g<sup>-1</sup>) | K<br>(mg g<sup>-1</sup>) | Ca<br>(mg g<sup>-1</sup>) | Mg<br>(mg g<sup>-1</sup>) | Na<br>(mg g<sup>-1</sup>) | Fe<br>(mg g<sup>-1</sup>) | Mn<br>(μg g<sup>-1</sup>) | Mo<br>(μg g<sup>-1</sup>) | Zn<br>(μg g<sup>-1</sup>) |
| --- | --- | --- | --- | --- | --- | --- | --- | --- | --- | --- | --- | --- | --- |
| *Biomass Harvesting* |  |  |  |  |  |  |  |  |  |  |  |  |  |
| SOH | 38.6±3.9 | 44±2 | 1.41±0.15 | 27.7±0.8 | 29.3±2.2 | 4.07±0.2 | 0.82±0.06 | 0.17±0.09 | 1.17±0.09 | 1.8±0.4 | 41.5±3.5 | 0.16±0.01 | 6.60±0.29 |
| AAH | 35.4±3.9 | 42±4 | 1.46±0.15 | 24.5±0.8 | 25.3±2.5 | 5.40±0.2 * | 0.98±0.06 | 0.25±0.09 | 1.65±0.08 * | 3.1±0.3 * | 62.0±3.5 * | 0.21±0.01 | 7.37±0.48 |
| BAH | 30.2±3.9 | 39±2 | 1.14±0.15 | 26.3±0.8 | 25.3±2.9 | 4.31±0.2 | 0.71±0.04 | 0.15±0.08 | 1.22±0.08 | 2.2±0.3 | 43.6±3.5 | 0.15±0.01 | 5.77±0.36 |
| WTH | 31.1±3.9 | 31±2 * | 1.10±0.15 | 28.2±0.8 | 21.0±3.5 | 4.09±0.2 | 0.72±0.03 | 0.11±0.04 | 1.11±0.05 | 1.5±0.3 | 35.4±3.5 | 0.13±0.01 | 3.85±0.54 |
| *Compensation Methods* |  |  |  |  |  |  |  |  |  |  |  |  |  |
| Control | 37.1±2.9 | 39±1 | 1.36±0.11 | 27.2±0.7 | 27.1±2.4 | 4.47±0.13 | 0.85±0.03 | 0.18±0.10 | 1.31±0.05 | 2.1±0.3 | 46.6±3.2 | 0.16±0.01 | 5.69±0.31 |
| Phosphorus | 33.6±2.9 | 42±2 | 1.28±0.11 | 26.3±0.7 | 24.5±2.4 | 4.45±0.26 | 0.74±0.03 * | 0.16±0.08 | 1.26±0.09 | 2.0±0.3 | 45.0±4.9 | 0.17±0.01 | 5.57±0.32 |
| Ash | 30.8±2.9 * | 36±1 * | 1.18±0.11 * | 26.5±0.7 | 24.0±2.4 | 4.50±0.15 | 0.85±0.03 | 0.17±0.09 | 1.31±0.06 | 2.3±0.3 | 45.2±2.0 | 0.16±0.01 | 6.42±0.36 |

All values are for the total content of the elements in the soil (except for P). SOC = soil organic carbon; DOC = dissolved organic carbon. For phosphorus (P), values are extractable P using the Olsen method. SOH = stem-only harvest; AAH = aboveground additional harvest; BAH = belowground additional harvest; WTH = whole-tree harvest. Control = no nutrient application. Treatments with an asterisk differ at $P < 0.05$ from their reference treatment (SOH or Control, respectively; see Methods). Mean values of the Biomass Harvesting treatments (n = 3 blocks) were calculated by pooling the Compensation Methods subplots. Mean values of the Compensation Methods treatments were calculated based on 12 replicates (3 blocks × 4 BH treatments).

**Table 3.** Soil acidity and exchangeable cations as affected by biomass harvest and compensation methods

|  | pH-H<sub>2</sub>O<br>unit less | pH-CaCl<sub>2</sub><br>unit less | CEC<sub>m</sub><br>(cmol.<sub>c</sub> kg<sup>-1</sup>) | BS<br>(%) | K<br>(cmol.<sub>c</sub> kg<sup>-1</sup>) | Ca<br>(cmol.<sub>c</sub> kg<sup>-1</sup>) | Mg<br>(cmol.<sub>c</sub> kg<sup>-1</sup>) | Na<br>(cmol.<sub>c</sub> kg<sup>-1</sup>) |
| --- | --- | --- | --- | --- | --- | --- | --- | --- |
| *Biomass Harvesting* |  |  |  |  |  |  |  |  |
| SOH | 4.43±0.07 | 3.43±0.08 | 3.51±0.23 | 43±2 | 0.06±0.00 | 0.92±0.04 | 0.36±0.02 | 0.06±0.00 |
| AAH | 4.68±0.07 | 3.69±0.08 | 2.90±0.24 | 38±2 | 0.06±0.00 | 0.67±0.03 * | 0.21±0.01 * | 0.05±0.00 |
| BAH | 4.49±0.07 | 3.57±0.08 | 2.51±0.23 * | 42±2 | 0.05±0.00 | 0.65±0.04 * | 0.23±0.02 * | 0.04±0.00 * |
| WTH | 4.51±0.07 | 3.46±0.08 | 2.90±0.24 | 39±2 | 0.06±0.01 | 0.69±0.06 | 0.28±0.02 * | 0.04±0.00 * |
| *Compensation Methods* |  |  |  |  |  |  |  |  |
| Control | 4.41±0.06 | 3.47±0.07 | 3.45±0.21 | 39±2 | 0.06±0.00 | 0.74±0.05 | 0.29±0.01 | 0.05±0.00 |
| Phosphorus | 4.53±0.05 | 3.51±0.07 | 2.58±0.19 * | 38±2 | 0.06±0.00 | 0.57±0.04 * | 0.22±0.02 * | 0.04±0.00 * |
| Ash | 4.65±0.05 * | 3.64±0.07 | 2.84±0.19 * | 46±2 * | 0.06±0.00 | 0.89±0.04 * | 0.30±0.02 | 0.05±0.00 |

SOH = stem-only harvest; AAH = aboveground additional harvest; BAH = belowground additional harvest; WTH = whole-tree harvest. Control = no nutrient application. CEC<sub>m</sub> = cationic exchange capacity (measured value); BS = base saturation of the CEC, which is the proportion of charges occupied by non-acidic exchangeable cations (K, Ca, Mg, Na). For consistency between the different terms of its equation, BS was calculated using the effective CEC (CEC<sub>e</sub>: calculated as the sum all the charges of all the analysed cations), whose values differ slightly from CEC<sub>m</sub> values. Treatments with an asterisk differ at $P < 0.05$ from their reference treatment (SOH or Control, respectively; see Methods). Mean values of the Biomass Harvesting treatments (n = 3 blocks) were calculated by pooling the Compensation Methods subplots. Mean values of the Compensation Methods treatments were calculated based on 12 replicates (3 blocks × 4 BH treatments).

**Table 4.**
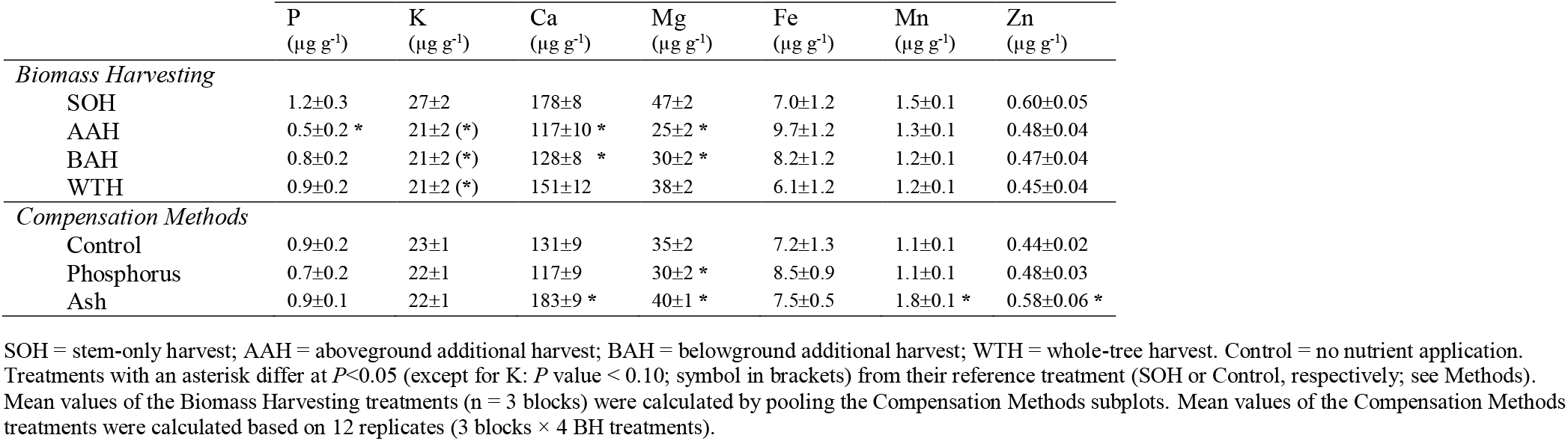
Soil composition in NH_4_-NO_3_ extractable nutrients and micronutrients as affected by biomass harvest and compensation methods

The effects of the treatments on P also depended on the chemical form considered. The soil P content as measured using the sodium bicarbonate extractant (P_-Olsen_) showed no significant differences, even if the non-conventional harvest treatments resulted in a decreasing trend in this soil content (Table 2). This not-significant trend became significant when considering the P content as measured using the NH_4_-NO_3_ extractant (Table 4). As expected, the application of wood ash did not significantly modify the soil P status. Surprisingly the P-fertiliser treatment did not improve this P status either.

The soil N_-total_ content followed the same, but insignificant, trend as for P with low values in the treatments that involved canopy harvesting (Table 2). The C/N ratio was remarkably irresponsive to all experimental treatments, but the soil N content was reduced by wood ash application.

### Vegetation response (H2)

The total aboveground biomass of the spontaneous vegetation remained unaffected by the *Biomass Harvesting* treatments, but changed due to the *Compensatory Methods* treatments with an increasing standing biomass following the ranking: Control ≤ P fertilisation ≤ Ash (Figure 2). For both factors, the proportion of the prominent plant species was influenced by treatments. For the *Biomass Harvesting* factor (BH), the proportion of species adapted to oligotrophic-acidic moorlands (*i.e. Molinia caerulea* and ericaceous species) was increased by the intensive harvests as compared with the conventional SOH treatment (Figure 2). The opposite pattern was observed in the wood ash treatment (*Compensation Methods* factor, CM), which strongly decreased the abundance of those moorland-adapted species. The wood ash application and, to a lesser extent, the P fertilisation treatments enhanced the growth of the N-fixers (almost entirely composed of the spiny shrub *Ulex europaeus*).

**Figure 2.**
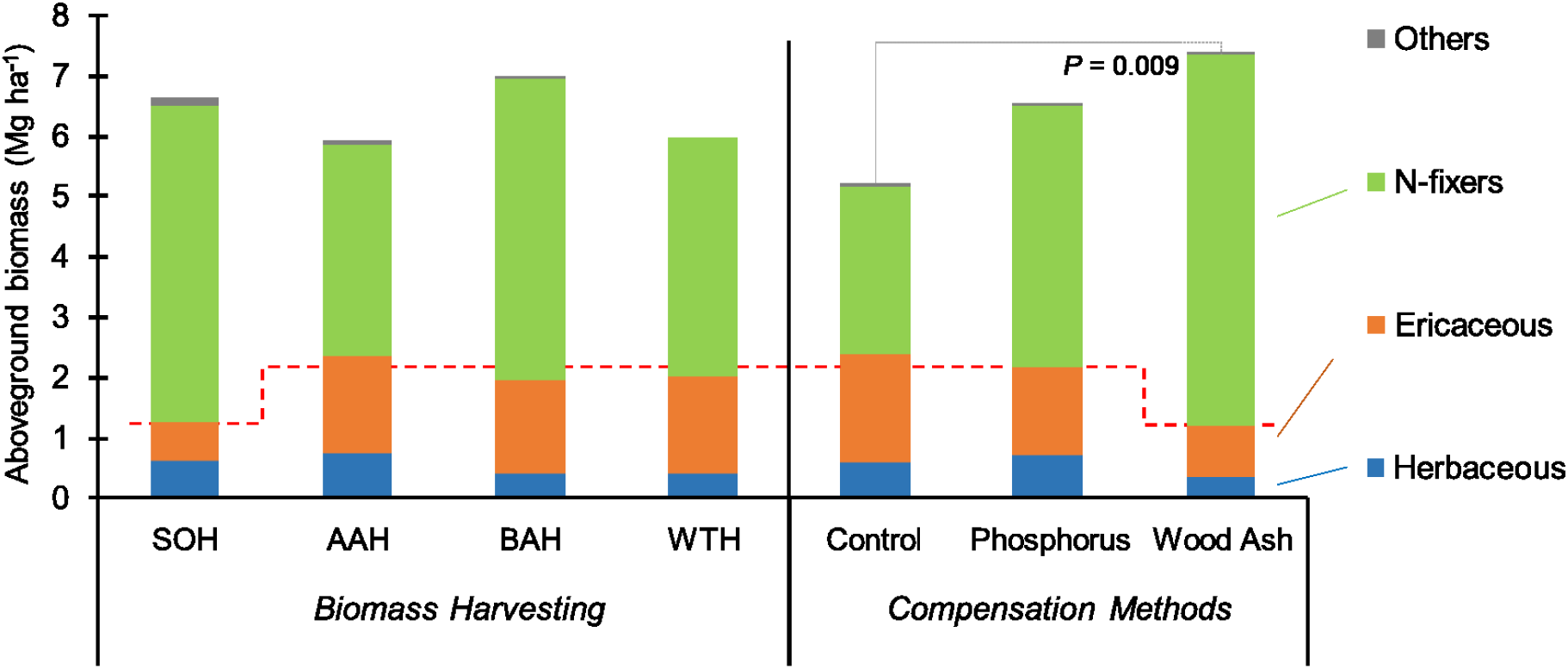
Understorey aboveground biomass as affected by biomass harvest and compensation methods. SOH = stem-only harvest; AAH = aboveground additional harvest; BAH = belowground additional harvest; WTH = whole-tree harvest. Control = no nutrient application. Mean values of the Biomass Harvesting treatments (n = 3 blocks) were calculated by pooling the Compensation Methods subplots. Mean values of the Compensation Methods treatments were calculated based on 12 replicates (3 blocks × 4 BH treatments). The understorey biomass was significantly higher in the Wood Ash treatment than in the Control treatment. The relative abundance of the species typical of oligotrophic-acidic conditions (herbaceous *Molinia caerula* and ericaceous *Calluna vulgaris* & *Erica scoparia*) is highlighted by the red dotted line. Photos of the main species are given in the Figure S4.

Measuring the trees at 7 years-old suggested that treatments did not affect the initial stand growth as no significant differences were observed (data not presented). At 11 years-old, the most severe treatments of the *Biomass Harvesting* factor (*i.e*. AAH and WTH) tended to depress the tree growth (field observations and Figure 3), but these trends remained statistically not-significant. Conversely we measured an unexpected significant difference between two treatments of the *Compensatory Methods* factor, with a negative effect of P fertilisation, and a positive trend of wood ash application (Figure 3; Table S3). The wood ash trend can be related to the improvement of the nutritional status of foliage (N, P, K, Mg, B, Cu, and Fe; Table 5).

**Figure 3.**
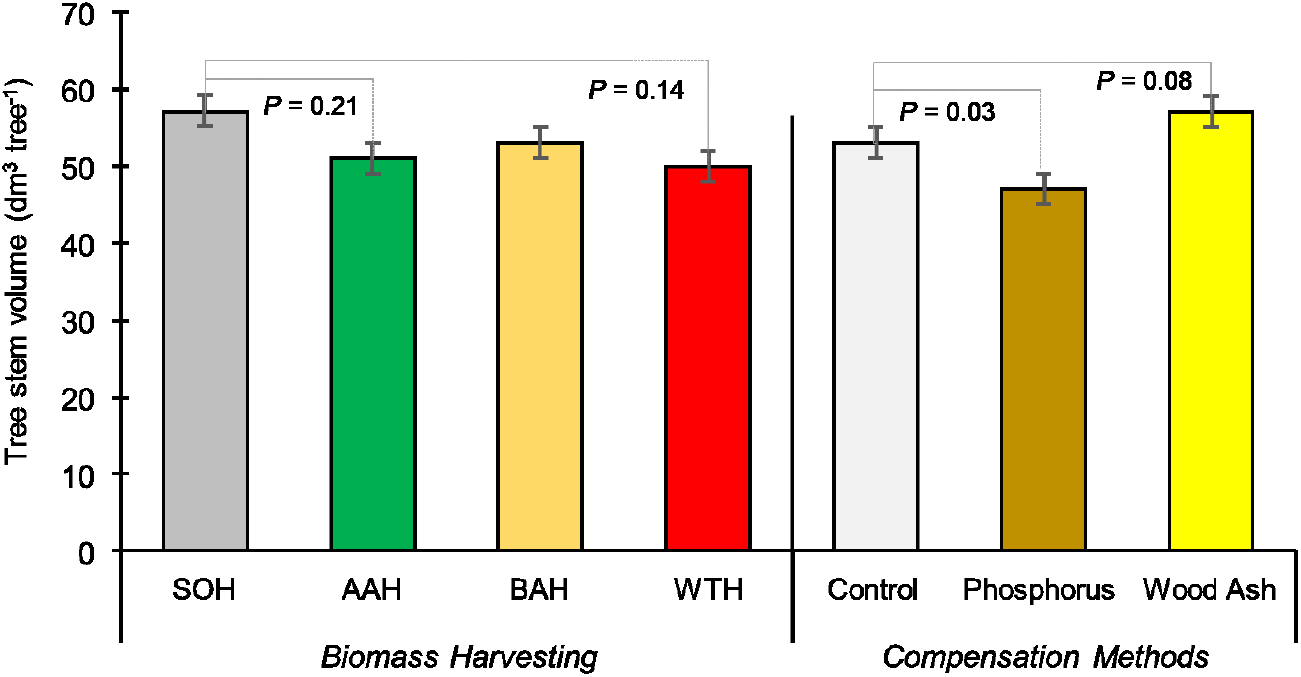
Tree stem volume as affected by biomass harvest and compensation methods. SOH = stem-only harvest; AAH = aboveground additional harvest; BAH = belowground additional harvest; WTH = whole-tree harvest. Control = no nutrient application. Error bars are standard errors. Mean values of the Biomass Harvesting treatments (n = 3 blocks) were calculated by pooling the Compensation Methods subplots. Mean values of the Compensation Methods treatments were calculated based on 12 replicates (3 blocks × 4 BH treatments). *P* values for differences between treatments of reference (*i.e*. SOH and Control) on the one hand, and other treatments on the other hand, are indicated in the graph. Trees were measured at 11 years old. Other tree metrics are shown in Table S3.

**Table 5.** Tree foliage composition (nutrients and micronutrients) as affected by biomass harvest and compensation methods

|  | N<br>(mg g<sup>-1</sup>) | P<br>(mg g<sup>-1</sup>) | K<br>(mg g<sup>-1</sup>) | Ca<br>(mg g<sup>-1</sup>) | Mg<br>(mg g<sup>-1</sup>) | B<br>(μg g<sup>-1</sup>) | Cu<br>(μg g<sup>-1</sup>) | Fe<br>(μg g<sup>-1</sup>) | Mn<br>(μg g<sup>-1</sup>) | Mo<br>(μg g<sup>-1</sup>) | Zn<br>(μg g<sup>-1</sup>) |
| --- | --- | --- | --- | --- | --- | --- | --- | --- | --- | --- | --- |
| *Biomass Harvesting* |  |  |  |  |  |  |  |  |  |  |  |
| SOH | 12.8±0.7 | 0.64±0.04 | 4.6±0.1 | 2.3±0.1 | 1.4±0.1 | 16.3±1.0 | 1.8±0.2 | 27.8±2.0 | 96.3±11.0 | 0.011±0.003 | 29.8±2.5 |
| AAH | 13.9±0.7 | 0.69±0.04 | 4.4±0.1 | 2.2±0.1 | 1.5±0.1 | 16.7±1.0 | 1.8±0.2 | 32.1±2.1 * | 104.2±10.9 | 0.014±0.002 | 32.1±2.5 |
| BAH | 12.1±0.7 | 0.65±0.04 | 4.5±0.1 | 2.2±0.1 | 1.4±0.1 | 17.2±1.0 | 2.0±0.2 | 27.1±2.0 | 92.9±11.7 | 0.009±0.003 | 30.5±2.5 |
| WTH | 12.6±0.7 | 0.65±0.05 | 4.6±0.1 | 2.3±0.1 | 1.4±0.1 | 17.2±1.0 | 1.8±0.3 | 28.9±2.9 | 81.0±11.7 | 0.012±0.002 | 32.1±2.5 |
| *Compensation Methods* |  |  |  |  |  |  |  |  |  |  |  |
| Control | 12.3±0.7 | 0.64±0.04 | 4.2±0.1 | 2.2±0.1 | 1.4±0.1 | 16.0±0.9 | 1.7±0.1 | 27.0±2.1 | 92.3±9.8 | 0.011±0.002 | 29.1±2.2 |
| Phosphorus | 13.6±0.7 * | 0.66±0.04 * | 4.5±0.1 * | 2.3±0.1 | 1.5±0.1 * | 18.0±0.9 * | 1.9±0.1 | 30.4±2.1 * | 100.1±9.4 * | 0.012±0.002 | 32.7±2.2 * |
| Ash | 12.7±0.7 * | 0.67±0.04 * | 4.8±0.1 * | 2.3±0.1 | 1.4±0.1 | 16.6±0.9 * | 2.0±0.1 | 29.6±2.0 * | 88.4±9.8 | 0.012±0.002 | 31.7±2.2 * |

SOH = stem-only harvest; AAH = aboveground additional harvest; BAH = belowground additional harvest; WTH = whole-tree harvest. Control = no nutrient application. Treatments with an asterisk differ at $P < 0.05$ from their reference treatment (SOH or Control, respectively; see Methods). Mean values of the Biomass Harvesting treatments (n = 3 blocks) were calculated by pooling the Compensation Methods subplots. Mean values of the Compensation Methods treatments were calculated based on 12 replicates (3 blocks × 4 BH treatments).

### Soil organic matter and CEC (H3)

The soil Cation Exchange Capacity (CEC) was strongly and linearly correlated with the quantity of elements that are commonly found as organic matter in acidic soils, which are total nitrogen (N_-total_) and organic carbon (SOC; CEC = 0.088 × SOC; r^2^ = 0.76). Due to the tight proportional relationship between CEC and SOC, the observed effects of the experimental treatments on these variables were quite similar. The harvest of the belowground biomass and the application of wood ash both tended to decrease the soil organic matter content and the soil CEC (Tables 2 and 3). The dissolved organic carbon (DOC) followed a similar pattern with a significant decrease in concentration after whole-tree harvesting (WTH) and wood ash application (Table 2).

### Soil acidity status (H4)

Harvesting more biomass than conventionally practiced did not modify any of the soil variables used to assess the acidic status of a soil (pH-H_2_O, pH-CaCl_2_, Base Saturation of the CEC; Table 3). The absence of effect on the Base Saturation was the consequence of the concomitant decrease in *basic cations* (K^+^, Ca^2+^, Mg^2+^, and Na^+^) and CEC. Conversely to biomass harvesting, applying wood ash modified the soil acidity status. The soil pH-H_2_O value increased by +0.24 unit, and the Base Saturation increased from 39% to 46% of the CEC (Table 3). The P fertilisation had no effect on soil acidity.

### Ecosystem content in trace metals (H5)

Overall, the *Biomass Harvesting* treatments generally had no significant effect on the trace elements (As, Cd, Mo, Ni, Pb, Tl, and Zn) in the ecosystem (Tables 2, 4, 5, 6, S4). The exceptions were a decrease in soil Pb and Zn contents due to WTH.

**Table 6.**
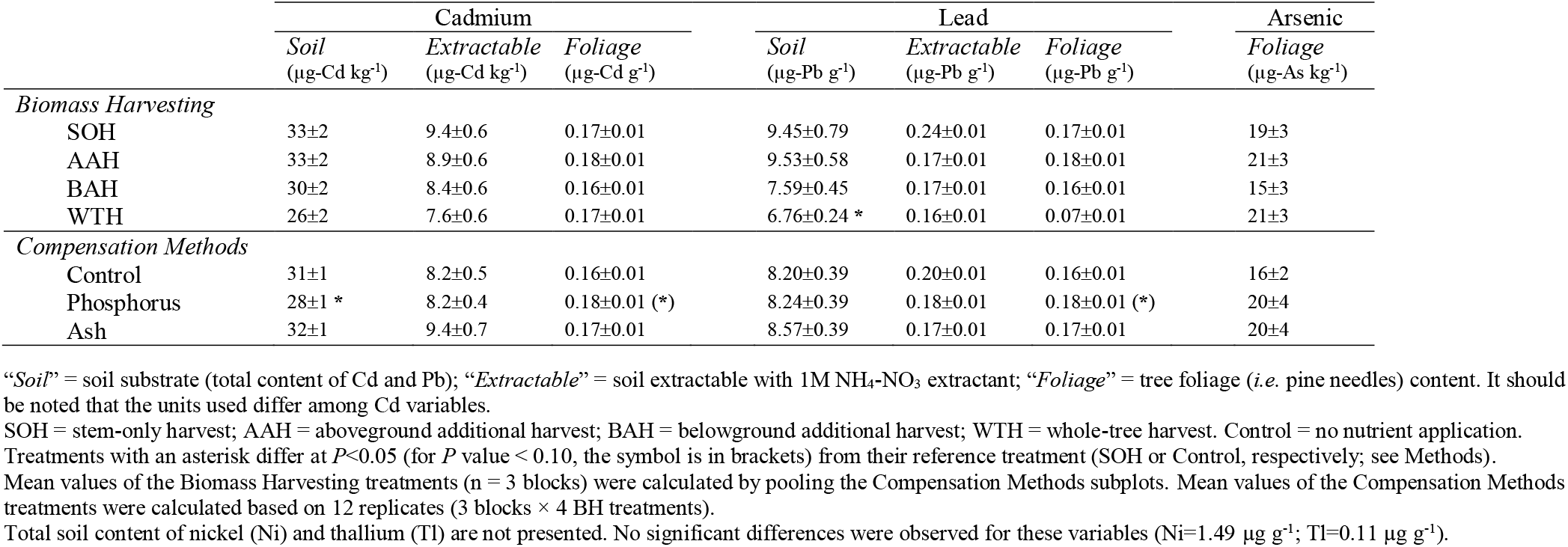
Ecosystem content in non-essential trace metals

The *Compensatory Methods* also had only a few significant effects on the trace element distribution in the ecosystem (Tables 2, 4, 5, 6, S4). The most noticeable exceptions were increases of soil Zn content due to wood ash application (Tables 4 and S4), an effect of P fertilisation on Cd distribution between the soil (decrease of Cd soil content) and trees (increase of Cd foliage content), and an improvement of Cu content of tree foliage after P or wood ash application (Table 6). This lack of effects can be explained by comparing the applied amounts with the initial soil stocks: for Fe, Mn, and Pb, the doses were 1-2 % of the initial stocks (0-15 cm soil layer). For Cd and Zn, the doses applied represented 8-10% of the initial stocks (Cd: 8% and 10% by fertilisation and wood ash, respectively; Zn: 8% by ash).

### Long term perspective: estimates of input-output budgets

The effects of treatments were highly nutrient-dependent. For nitrogen, the budget values were positive or close to equilibrium for the SOH, but were negative at 11 years for the other levels of biomass harvest (Figure S5). The budget values became positive over time, mainly due to atmospheric deposition and nitrogen symbiotic fixation. Applying P-fertiliser or wood ash directly improved budgets by enhancing the growth of the spontaneous N-fixers (Figure 2). In the case of phosphorus, because most external sources of P were negligible (*i.e*. < 1 kg ha^-1^ yr^-1^), only applying P-fertiliser enabled to balance the biomass harvests.

The BAH and WTH treatments had a strong impact on K input-output budgets because the belowground biomass of trees was particularly rich in this nutrient. In these cases, applying wood ash and keeping long silvicultural rotations enabled to balance the exports (Figure S5). The same pattern was observed for calcium and magnesium but to different extents, depending on the biomass nutrient content and the compensatory fluxes that are atmospheric deposition and mineral weathering.

## Discussion

### Importance of the study design and the study region for data interpretation

Implementing a careful harvest of adult tree compartments over small areas was hardly feasible because the machines that collect stumps or canopies need space to work (Figure S3). Consequently, the *Biomass Harvesting* factor could be studied only by using large plots (*i.e*. one ha per plot). With 16 experimental treatments and three blocks, building a complete factorial experimental design (*i.e*. with 48 large plots) was not possible because the surface area of 48 ha required was incompatible with the premise of an area that was initially homogeneous in its properties and past-management. Therefore, we built our experiment based on a split-plot design. Such a design had one major consequence on data analysis. Indeed, the number of the subplots −dedicated to the *Compensation Methods* treatments− were four-fold more numerous than the experimental areas dedicated to *Biomass Harvesting* treatments (Figure S1). The power of the statistical tests was consequently much lower for the *Biomass Harvesting* factor, so isolating significant effects was sometimes difficult.

The forest region where our experiment took place is atypical from a biogeochemical perspective as the soils are extremely poor in phosphorus. While in natural systems the soil P-total content ranges between ~10 and 2,000 μg g^-1^ (Achat et al. 2009; Yang and Post 2011; Augusto et al. 2017), the study region has a mean value of ~50 μg g^-1^ and is within a range of ~10-100 μg g^-1^ (Achat et al. 2009; Augusto et al. 2010). Being also acidic and poor in other nutrients and micronutrients (Augusto et al. 2010; Trichet et al. 2018), the soils of the study region have been identified as having the highest level of vulnerability to biomass exports according to the national system of forest evaluation (Durante et al. 2019). Poorest of the poor, the study region is also expected to respond strongly and negatively to additional harvests of biomass, which implies that results should be extrapolated to other regions with caution.

### Consequences of harvesting more biomass on the biogeochemistry of an oligotrophic forest

Exporting tree canopies generally enhances the amount of harvested biomass (~ +20% to +50%), but this is at the expense of losing huge quantities of nutrients (~ +100% to +250%; (Achat et al. 2015a)). Our results (10-34% extra biomass, and 37-145% extra nutrients) fit well with this general pattern. On the other hand, the quantities of exported nutrients of this field experiment are about half of the values used in previous modelling studies (Augusto et al. 2015b; Achat et al. 2018). This was the direct consequences of the harvesting techniques that were purposely used to reduce the losses of nutrients: letting branches shed their needles (Stupak et al. 2008), and harvesting only stumps and coarse roots (Augusto et al. 2015a). Despite these precautions, harvesting more biomass, and above all harvesting the whole-tree, reduced the soluble pools of nutrients (K, Ca, Mg, and P) in the soil (−15% to −58%; Table 4). Additional biomass harvests likely also impacted the soil content in organic matter, with negative ripple effects on organic P and CEC, the latter being entirely dependent on organic matter in these sandy soils (Augusto et al. 2010). In the study region the organic forms of P are important for seedling and tree nutrition (Jonard et al. 2009; Achat et al. 2013), which may explain why we observed a depressive −but not significant− effect of harvesting canopies on tree growth in the subsequent stand (see Table S3). A longer monitoring is needed to confirm the observed trend. The absence of change in the tree foliage composition might be considered as surprising as foliage nutrient content is often used to assess plant nutrition (CSIRO 1997; Mellert and Göttlein 2012). Nevertheless, maintaining the nutrient content of foliage at the expense of growth is a response commonly observed after the harvest of additional biomass (Achat et al. 2015a), and our results are consistent with this general pattern.

### Fertilisation and wood ash application as possible mitigation practices

The major effect of applying wood ash was expected to be a change in the acidity status of soils (Reid and Watmough 2014). Our experiment was not an exception and the soil pH, base saturation and concomitant available calcium and magnesium, were improved by wood ash application (Gomez-Rey et al. 2013). The available soil micronutrient contents of Mn and Zn also logically increased because wood ash contains them in substantial quantities (Table S1).

Whereas the effects of wood ash were expected and straight forward to interpret, the P fertilisation had surprising outcomes: the soil P status was not improved, and the soil contents of available Ca and Mg actually decreased by P fertilisation. These results may be explained by the tree foliage composition that was modified by both P fertilisation and wood ash application, with increases in concentration values for most nutrients and micronutrients (and even some non-essential trace metals). Hence, it seemed that the two compensatory methods improved tree nutrition but, for P fertilisation, it was at the expense of the soil reserves in Ca and Mg.

In turn, these changes in the soil properties modified the dynamics of the spontaneous vegetation and of the tree plantation. The vegetation experienced a shift of species prominence after the wood ash application, with a decrease in the species typical of acidic moorlands and heathlands (*i.e. Molinia caerulea* and ericaceous species) to the benefit of the local main N-fixer (the shrub *Ulex europaeus*). The decrease of the abundance of the acidophilus species is logical since wood ash decreased the soil acidity. The increase of the N-fixer shrub was expected as we are used to observing a positive response of this leguminous species to P fertilisation (*e.g*. Delerue et al. (2015)), but although the N-fixers abundance was increased by P fertilisation, it was relatively low compared with previous experiments (Augusto et al. 2005; Vidal et al. 2019), and lower than results after the wood ash application. This ash effect may be the positive consequence of reducing the ambient acidity on the symbionts of the leguminous species (Slattery and Coventry 1995).

Trees that were planted responded differently to the *Compensatory Methods*. Firstly, trees that received wood ash tended to grow the fastest. Since wood ash application improved the availability of many nutrients and micronutrients, it suggests that plant growth in this site was limited by several elements at the same time. On the other hand, the depressive effect of P fertilisation was surprising considering that this practice has proven its efficiency in the study region for more than half a century (Trichet et al. 2009). As explained above, it seems that P fertilisation improved the tree foliage composition at the expense of soil fertility, and we speculate that this soil impoverishment might be at the origin of subsequent degraded tree growth. However, this atypical phenomenon requires further investigation. Considering the positive effects of wood ash application on soil acidity and nutrient content, and its positive repercussion on tree growth, one might conclude that this compensatory method should be promoted in oligotrophic forests submitted to intensive biomass harvestings. Nevertheless, this conclusion does not take into account one major drawback of wood ash application in our experiment, which was a decrease of the soil organic matter content, with negative consequences on organic carbon and total nitrogen contents and soil exchange capacity (losses ≈ 7-11%, after correcting for the change in soil bulk density). We interpret this decrease as the result of the acidity alleviation by wood ash, which probably enhanced the soil microbial activity and respiration (Bååth and Arnebrant 1994; Jokinen et al. 2006; Omil et al. 2013). In a meta-analysis, we previously concluded that wood ash had no influence on soil organic carbon (Augusto et al. 2008a), but this study was almost entirely based on Nordic, or cold temperate experiments (mean annual temperature < 8.5°C; *e.g*. Feldkirchner et al. (2003)). Under a warm temperate climate, wood ash application in sites with a long history of forest occupation (*i. e*. high SOC values and often low pH values) can decrease the soil content in organic carbon (Solla-Gullon et al. 2006).

### Initial expectations and future anticipation

Most, but not all, of our initial expectations were confirmed by the field experiment. Firstly, as anticipated, harvesting more biomass aggravated the P scarcity of the local soils, and the application of wood ash had only a minor influence on the soil P content (Hypothesis H1; Table S5). Still in line with our expectations, the impoverishment of the soil caused by additional harvests of biomass could result in reduced growth of the subsequent forest stand; wood ash application had positive effects on early tree growth (H2). The fact that, unexpectedly, P application did not enable trees to overcome P limitation suggested that other nutritional limitations −such as Ca and Mg− prevented the experimental fertilisation from having an effect. If our two first hypotheses were fairly well supported by results, the third and fourth hypotheses received mixed support. Indeed, both high rates of biomass harvesting and wood ash application decreased the pool of soil organic carbon, but only the former treatment was supposed to do so (H3). Similarly, while wood ash application logically improved the acidity status of the soil to a moderate extent, intensive biomass exports did not deteriorate it, as in many other studies (H4; see above for possible explanations). Finally, as expected, none of the experimental factors had major effects on the trace metal distribution within the ecosystem (H5).

Overall, this factorial experiment showed that exporting more forest biomass though the additional harvests of tree canopies, stumps, and roots had negative consequences on soil properties. Additional harvests have aggravated the poverty of the already oligotrophic soil, which in turn may decrease tree growth and the soil content of organic carbon in the future. Importantly, applying nutrients as fertiliser or wood ash did not fully compensate for the negative impact of biomass exports. Indeed, our estimates of input-output budgets suggests that, in addition to applying compensatory treatments, maintaining quite long silvicultural rotations is more suitable to ensure the sustainability of the forest management.

## Supporting information

Mean values per treatment

## Acknowledgements

We are grateful to the ADEME, GIS G-PMF and the Council of the Nouvelle-Aquitaine region for their financial support (D.V. PhD grant; SYLVOGENE and PINASTER projects). Similarly, we thank the ISPA lab for providing the F.B. Master grant. We are grateful to the Forest Experimental Facility (UEFP-https://doi.org/10.15454/1.5483264699193726E12) for their assistance during the successive field campaigns. We thank Sylvie Bussière, Cécile Coriou, and Catherine Lambrot for their help during the sample analyses. Finally, we thank Jean-Luc Denou and Pierre Bordenave for providing data from the XyloSylve monitoring experiment.

## References

Achat DL, Bakker MR, Augusto L, et al (2009) Evaluation of the phosphorus status of P-deficient podzols in temperate pine stands: combining isotopic dilution and extraction methods. Biogeochemistry 92:183–200. doi: 10.1007/s10533-008-9283-7

Achat DL, Bakker MR, Augusto L, Morel C (2013) Contributions of microbial and physical-chemical processes to phosphorus availability in Podzols and Arenosols under a temperate forest. Geoderma 211:18–27. doi: DOI 10.1016/j.geoderma.2013.07.003

Achat DL, Deleuze C, Landmann G, et al (2015a) Quantifying consequences of removing harvesting residues on forest soils and tree growth–A meta-analysis. For Ecol Manage 348:124–141. doi: doi: 10.1016/j.foreco.2015.03.042

Achat DL, Fortin M, Landmann G, et al (2015b) Forest soil carbon is threatened by intensive biomass harvesting. Sci Rep 5:15991

Achat DL, Martel S, Picart D, et al (2018) Modelling the nutrient cost of biomass harvesting under different silvicultural and climate scenarios in production forests. For Ecol Manage 429:642–653. doi: 10.1016/j.foreco.2018.06.047

Alvarez-Alvarez P, Pizarro C, Barrio-Anta M, et al (2018) Evaluation of tree species for biomass energy production in Northwest Spain. FORESTS 9:. doi: 10.3390/f9040160

Andre F, Jonard M, Ponette Q (2010) Biomass and nutrient content of sessile oak (Quercus petraea (Matt.) Liebl.) and beech (Fagus sylvatica L.) stem and branches in a mixed stand in southern Belgium. Sci Total Environ 408:2285–2294. doi: DOI 10.1016/j.scitotenv.2010.02.040

Aronsson KA, Ekelund NGA (2004) Biological effects of wood ash application to forest and aquatic ecosystems. J Environ Qual 33:1595–1605. doi: 10.2134/jeq2004.1595

Augusto L, Achat DL, Bakker MR, et al (2015a) Biomass and nutrients in tree root systems–sustainable harvesting of an intensively managed Pinus pinaster (Ait.) planted forest. Glob Chang Biol Bioenergy 7:231–243. doi: doi: 10.1111/gcbb.12127

Augusto L, Achat DL, Jonard M, et al (2017) Soil parent material - A major driver of plant nutrient limitations in terrestrial ecosystems. Glob Chang Biol 23:3808–3824

Augusto L, Bakker MR, De Lavaissiere C, et al (2009) Estimation of nutrient content of woody plants using allometric relationships: quantifying the difference between concentration values from the literature and actuals. Forestry 82:463–477. doi: 10.1093/forestry/cpp019

Augusto L, Bakker MR, Meredieu C (2008a) Wood ash applications to temperate forest ecosystems - potential benefits and drawbacks. Plant Soil 306:181–198. doi: 10.1007/s11104-008-9570-z

Augusto L, Bakker MR, Morel C, et al (2010) Is “grey literature” a reliable source of data to characterize soils at the scale of a region? A case study in a maritime pine forest in southwestern France. Eur J Soil Sci 61:807–822

Augusto L, Crampon N, Saur E, et al (2005) High rates of nitrogen fixation of Ulex species in the understory of maritime pine stands and the potential effect of phosphorus fertilization. Can J For Res Can Rech For 35:1183–1192. doi: 10.1139/x05-054

Augusto L, De Schrijver A, Vesterdal L, et al (2015b) Influences of evergreen gymnosperm and deciduous angiosperm tree species on the functioning of temperate and boreal forests. Biol Rev 90:444–466

Augusto L, Meredieu C, Bert D, et al (2008b) Improving models of forest nutrient export with equations that predict the nutrient concentration of tree compartments. Ann For Sci 65:. doi: 10.1051/forest:2008059

Bååth E, Arnebrant K (1994) Growth rate and response of bacterial communities to pH in limed and ash treated forest soils. Soil Biol Biochem 26:995–1001

Banos V, Dehez J (2017) Le bois-énergie dans la tempête, entre innovation et captation ? Les nouvelles ressources de la forêt landaise. Natures Sci Sociétés 25:122–133

Bertran P, Bateman MD, Hernandez M, et al (2011) Inland aeolian deposits of south-west France: facies, stratigraphy and chronology. J Quat Sci 26:374–388

Bredoire F, Bakker MR, Augusto L, et al (2016) What is the P value of Siberian soils? Soil phosphorus status in south-western Siberia and comparison with a global data set. Biogeosciences 13:2493–2509. doi: 10.5194/bg-13-2493-2016

Bright RM, Cherubini F, Astrup R, et al (2012) A comment to “Large-scale bioenergy from additional harvest of forest biomass is neither sustainable nor greenhouse gas neutral”: Important insights beyond greenhouse gas accounting. Glob Chang Biol Bioenergy 4:617–619

Cavard X, Augusto L, Saur E, Trichet P (2007) Field effect of P fertilization on N2 fixation rate of Ulex europaeus. Ann For Sci 64:875–881. doi: 10.1051/forest:2007066

Croise L, Ulrich E, Duplat P, Jaquet O (2005) Two independent methods for mapping bulk deposition in France. Atmos Environ 39:3923–3941

CSIRO (1997) Plant analysis: an interpretation manual. CSIRO Publishing, Reuter, D. J. and Robinson, J. B. (Eds), Collingwood, Australia

de Schrijver A, de Frenne P, Staelens J, et al (2012) Tree species traits cause divergence in soil acidification during four decades of postagricultural forest development. Glob Chang Biol 18:1127–1140

Deirmendjian L, Loustau D, Augusto L, et al (2018) Hydro-ecological controls on dissolved carbon dynamics in groundwater and export to streams in a temperate pine forest. Biogeosciences 15:669–691

Delerue F, Gonzalez M, Michalet R, et al (2015) Weak Evidence of Regeneration Habitat but Strong Evidence of Regeneration Niche for a Leguminous Shrub. PLoS One 10:. doi: 10.1371/journal.pone.0130886

Durante S, Augusto L, Achat DL, et al (2019) Diagnosis of forest soil sensitivity to harvesting residues removal - A transfer study of soil science knowledge to forestry practitioners. Ecol Indic 104:512–523. doi: 10.1016/j.ecolind.2019.05.035

Feldkirchner DC, Wang C, Gower ST, et al (2003) Effects of nutrient and paper mill biosolids amendments on the growth and nutrient status of hardwood forests. For Ecol Manage 177:95–116. doi: 10.1016/S0378-1127(02)00318-3

Forrester DI, Pares A, O’hara C, et al (2013) Soil organic carbon is increased in mixed-species plantations of Eucalyptus and nitrogen-fixing Acacia. Ecosystems 16:123–132

Fransson AM, Bergkvist B, Tyler G (1999) Phosphorus solubility in an acid forest soil as influenced by form of applied phosphorus and liming. Scand J For Res 14:538–544. doi: 10.1080/02827589950154014

Garcia W de O, Amann T, Hartmann J (2018) Increasing biomass demand enlarges negative forest nutrient budget areas in wood export regions. Sci Rep 8:. doi: 10.1038/s41598-018-22728-5

Gomez-Rey MX, Madeira M, Coutinho J (2013) Soil C and N dynamics, nutrient leaching and fertility in a pine plantation amended with wood ash under Mediterranean climate. Eur J For Res 132:281–295. doi: 10.1007/s10342-012-0674-x

Gonzalez M, Augusto L, Gallet-Budynek A, et al (2013) Contribution of understory species to total ecosystem aboveground and belowground biomass in temperate Pinus pinaster Ait. forests. For Ecol Manage 289:38–47. doi: DOI 10.1016/j.foreco.2012.10.026

Haberl H, Schulze ED, Korner C, et al (2013) Response: complexities of sustainable forest use. Glob Chang Biol Bioenergy 5:1–2

Hagerberg D, Wallander H (2002) The impact of forest residue removal and wood ash amendment on the growth of the ectomycorrhizal external mycelium. FEMS Microbiol Ecol 39:139–146. doi: 10.1016/S0168-6496(01)00207-0

Hannam KD, Venier L, Allen D, et al (2018) Wood ash as a soil amendment in Canadian forests: what are the barriers to utilization? Can J For Res 48:442–450. doi: 10.1139/cjfr-2017-0351

IPCC (2014) Climate Change 2014: Synthesis Report. Fifth Assessment Report. Geneva, Switzerland

Jokinen H, Kiikkilä O, Fritze H (2006) Exploring the mechanisms behind elevated microbial activity after wood ash application. Soil Biol Biochem 38:2285–2291

Jonard M, Augusto L, Morel C, et al (2009) Forest floor contribution to phosphorus nutrition: experimental data. Ann For Sci 66:510. doi: 10.1051/forest/2009039

Jurevics A, Peichl M, Olsson BA, et al (2016) Slash and stump harvest have no general impact on soil and tree biomass C pools after 32-39 years. For Ecol Manage 371:33–41. doi: 10.1016/j.foreco.2016.01.008

Kabata-Pendias A (2000) Trace elements in soils and plants. CRC Taylor and Francis Group, London NewYork

Kimmins JP (1974) Sustained yield, timber mining, and the concept of ecological rotation, a British Columbian view. For Chron Feb 1974:27–31

Lindner M, Karjalainen T (2007) Carbon inventory methods and carbon mitigation potentials of forests in Europe: a short review of recent progress. Eur J For Res 126:149–156

Luyssaert S, Marie G, Valade A, et al (2018) Trade-offs in using European forests to meet climate objectives. Nature 562:259–262

Mellert KH, Göttlein A (2012) Comparison of new foliar nutrient thresholds derived from van den Burg’s literature compilation with established central European references. Eur J For Res 131:1461–1472

Mora O, Banos V, Regolini M, Carnus JM (2014) Using scenarios for forest adaptation to climate change: a foresight study of the Landes de Gascogne Forest 2050. Ann For Sci 71:313–324

Nicholls D, Monserud RA, Dykstra DP (2009) International bioenergy synthesis-Lessons learned and opportunities for the western United States. For Ecol Manage 257:1647–1655

Nnadi EO, Mbah CN, Nweke AI, Njoku C (2019) Physicochemical properties of an acid ultisol subjected to different tillage practices and wood-ash amendment: Impact on heavy metal concentrations in soil and Castor plant. Soil Tillage Res 194:104288

Nohrstedt HO (2001) Response of coniferous forest ecosystems on mineral soils to nutrient additions: A review of Swedish experiences. Scand J For Res 16:555–573. doi: 10.1080/02827580152699385

Omil B, Pineiro V, Merino A (2013) Soil and tree responses to the application of wood ash containing charcoal in two soils with contrasting properties. For Ecol Manage 295:199–212. doi: 10.1016/j.foreco.2013.01.024

Pitman RM (2006) Wood ash use in forestry - a review of the environmental impacts. FORESTRY 79:563–588. doi: 10.1093/forestry/cpl041

R Core Team (2019) R: A language and environment for statistical computing. R Foundation for Statistical Computing; Vienna; Austria. URL https://www.R-project.org

Ranger J, Turpault MP (1999) Input-output nutrient budgets as a diagnostic tool for sustainable forest management. For Ecol Manage 122:139–154

Ranius T, Hamalainen A, Egnell G, et al (2018) The effects of logging residue extraction for energy on ecosystem services and biodiversity: A synthesis. J Environ Manage 209:409–425. doi: 10.1016/j.jenvman.2017.12.048

Reid C, Watmough SA (2014) Evaluating the effects of liming and wood-ash treatment on forest ecosystems through systematic meta-analysis. Can J For Res 44:867–885. doi: 10.1139/cjfr-2013-0488

Saur E, Ranger J, Lemoine B, Gelpe J (1992) Micronutrient Distribution in 16-Year-Old Maritime Pine. Tree Physiol 10:307–316

Schulze ED, Korner CI, Law BE, et al (2012) Large-scale bioenergy from additional harvest of forest biomass is neither sustainable nor greenhouse gas neutral. Glob Chang Biol Bioenergy 4:611–616

Slattery JF, Coventry DR (1995) Acid-tolerance and symbiotic effectiveness of Rhizobium-Leguminosarum by Trifoloo isolated from subterranean clover growing in permanent pastures. Soil Biol Biochem 27:111–115. doi: 10.1016/0038-0717(94)00143-O

Solla-Gullon F, Santalla M, Rodriguez-Soalleiro RJ, Merino A (2006) Nutritional status and growth of a young Pseudotsuga menziesii plantation in a temperate region after application of wood-bark ash. For Ecol Manage 237:312–321. doi: 10.1016/j.foreco.2006.09.054

Steenari BM, Lindqvist O (1997) Stabilisation of biofuel ashes for recycling to forest soil. Biomass Bioenergy 13:39–50. doi: 10.1016/S0961-9534(97)00024-X

Stromgren M, Egnell G, Olsson BA (2013) Carbon stocks in four forest stands in Sweden 25 years after harvesting of slash and stumps. For Ecol Manage 290:59–66. doi: 10.1016/j.foreco.2012.06.052

Stupak I, Nordfjell T, Gundersen P (2008) Comparing biomass and nutrient removals of stems and fresh and predried whole trees in thinnings in two Norway spruce experiments. Can J For Res Can Rech For 38:2660–2673

Sverdrup H, Thelin G, Robles M, et al (2006) Assesing nutrient sustainability of forest production for different tree species considering Ca, Mg, K, N and P at Bjornstorp Estate, Sweden. Biogeochemistry 81:219–238

Symeonides C, McRae SG (1977) The assessment of plant-available cadmium in soils. J Environ Qual 6:120–123

Thiffault E, Hannam KD, Pare D, et al (2011) Effects of forest biomass harvesting on soil productivity in boreal and temperate forests - A review. Environ Rev 19:278–309

Trichet P, Bakker MR, Augusto L, et al (2009) Fifty years of fertilization experiments on Pinus pinaster in southwest France: the importance of phosphorus as a fertilizer. For Sci 55:390–402

Trichet P, Cheval N, Lambrot C, et al (2018) Using a dune forest as a filtering ecosystem for water produced by a treatment plant - One decade of environmental assessment. Sci Total Environ 640:849–861. doi: 10.1016/j.scitotenv.2018.05.263

Vance ED (1996) Land application of wood-fired and combination boiler ashes: An overview. J Environ Qual 25:937–944. doi: 10.2134/jeq1996.00472425002500050002x

Ventura M, Panzacchi P, Muzzi E, et al (2019) Carbon balance and soil carbon input in a poplar short rotation coppice plantation as affected by nitrogen and wood ash application. NEW For 50:969–990. doi: 10.1007/s11056-019-09709-w

Vidal DF, Trichet P, Puzos L, et al (2019) Intercropping N-fxing shrubs in pine plantation forestry as an ecologically sustainable management option. For Ecol Manage 437:175–187

Wall A (2012) Risk analysis of effects of whole-tree harvesting on site productivity. For Ecol Manage 282:175–184

Walmsley JD, Godbold DL (2010) Stump harvesting for bioenergy - A review of the environmental impacts. Forestry 83:17–38

Wang P, Olsson BA, Arvidsson H, Lundkvist H (2010) Short-term effects of nutrient compensation following whole-tree harvesting on soil and soil water chemistry in a young Norway spruce stand. Plant Soil 336:323–336. doi: 10.1007/s11104-010-0484-1

Yang X, Post WM (2011) Phosphorus transformations as a function of pedogenesis: A synthesis of soil phosphorus data using Hedley fractionation method. Biogeosciences 8:2907–2916

Zabowski D, Chambreau D, Rotramel N, Thies WG (2008) Long-term effects of stump removal to control root rot on forest soil bulk density, soil carbon and nitrogen content. For Ecol Manage 255:720–727

