## Supplementary material for "Response of soil and vegetation in a warm-temperate Pine forest to intensive biomass harvests, phosphorus fertilisation, and wood ash application": Mean values per treatment

**Appendix 1** – Mean values of the main responses variables of the study, presented in all factorial combinations of experimental factors

| Hypothesis | Variable | Biomass Harvesting (BH) | Compensation Methods (CM) | Mean value | Standard Error | Lower confidence limit | Upper confidence limit |
|---|---|---|---|---|---|---|---|
| H1 | P-Olsen | SOH | Control | 0.0070 | 0.0006 | 0.0044 | 0.0096 |
| *Soil fertility* | (g/100g) | | Ash | 0.0067 | 0.0006 | 0.0041 | 0.0093 |
| | | | P-fertiliser | 0.0063 | 0.0006 | 0.0037 | 0.0089 |
| | | AAH | Control | 0.0063 | 0.0007 | 0.0032 | 0.0095 |
| | | | Ash | 0.0048 | 0.0007 | 0.0017 | 0.0080 |
| | | | P-fertiliser | 0.0063 | 0.0007 | 0.0032 | 0.0095 |
| | | BAH | Control | 0.0067 | 0.0010 | 0.0025 | 0.0108 |
| | | | Ash | 0.0060 | 0.0010 | 0.0019 | 0.0101 |
| | | | P-fertiliser | 0.0048 | 0.0010 | 0.0007 | 0.0090 |
| | | WTH | Control | 0.0048 | 0.0013 | -0.0008 | 0.0104 |
| | | | Ash | 0.0040 | 0.0013 | -0.0016 | 0.0096 |
| | | | P-fertiliser | 0.0055 | 0.0013 | -0.0001 | 0.0111 |

| Hypothesis | Variable | Biomass Harvesting (BH) | Compensation Methods (CM) | Mean value | Standard Error | Lower confidence limit | Upper confidence limit |
|---|---|---|---|---|---|---|---|
| H2 | Tree height | SOH | Control | 87.5 | 1.9 | 79.5 | 95.4 |
| *Tree growth* | (dm) | | Ash | 88.4 | 1.9 | 80.4 | 96.4 |
| | | | P-fertiliser | 81.3 | 1.9 | 73.3 | 89.3 |
| | | AAH | Control | 79.9 | 1.9 | 71.9 | 87.9 |
| | | | Ash | 84.1 | 1.9 | 76.2 | 92.1 |
| | | | P-fertiliser | 82.8 | 1.9 | 74.8 | 90.8 |
| | | BAH | Control | 84.9 | 1.9 | 76.9 | 92.9 |
| | | | Ash | 87.0 | 1.9 | 79.0 | 95.0 |
| | | | P-fertiliser | 79.7 | 1.9 | 71.7 | 87.7 |
| | | WTH | Control | 84.4 | 1.9 | 76.4 | 92.4 |
| | | | Ash | 85.4 | 1.9 | 77.4 | 93.4 |
| | | | P-fertiliser | 82.0 | 1.9 | 74.0 | 90.0 |

| Hypothesis | Variable | Biomass Harvesting (BH) | Compensation Methods (CM) | Mean value | Standard Error | Lower confidence limit | Upper confidence limit |
|---|---|---|---|---|---|---|---|
| H3 | C-org | SOH | Control | 46.0 | 4.6 | 26.4 | 65.6 |
| *Soil* | (g/kg) | | Ash | 33.4 | 4.6 | 13.8 | 53.0 |
| *Organic* | | | P-fertiliser | 36.4 | 4.6 | 16.8 | 56.0 |
| *Matter* | | AAH | Control | 36.3 | 4.1 | 18.7 | 53.9 |
| | | | Ash | 33.3 | 4.1 | 15.7 | 50.9 |
| | | | P-fertiliser | 36.5 | 4.1 | 18.9 | 54.1 |
| | | BAH | Control | 35.8 | 3.9 | 19.0 | 52.7 |
| | | | Ash | 28.2 | 3.9 | 11.3 | 45.1 |
| | | | P-fertiliser | 26.4 | 3.9 | 9.6 | 43.3 |
| | | WTH | Control | 30.2 | 6.6 | 1.8 | 58.6 |
| | | | Ash | 28.3 | 6.6 | -0.1 | 56.7 |
| | | | P-fertiliser | 34.9 | 6.6 | 6.5 | 63.3 |

| Hypothesis | Variable | Biomass Harvesting (BH) | Compensation Methods (CM) | Mean value | Standard Error | Lower confidence limit | Upper confidence limit |
|---|---|---|---|---|---|---|---|
| H3 *Soil Organic Matter* | DOC (mg-C/L) | SOH | Control | 44.1 | 1.7 | 36.9 | 51.3 |
| | | | Ash | 41.5 | 4.5 | 22.0 | 61.0 |
| | | | P-fertiliser | 46.4 | 3.1 | 33.2 | 59.5 |
| | | AAH | Control | 39.2 | 4.4 | 20.2 | 58.2 |
| | | | Ash | 41.4 | 5.7 | 16.8 | 66.0 |
| | | | P-fertiliser | 52.8 | 11.6 | 3.1 | 102.5 |
| | | BAH | Control | 37.6 | 2.0 | 28.9 | 46.3 |
| | | | Ash | 35.6 | 1.7 | 28.4 | 42.8 |
| | | | P-fertiliser | 41.8 | 7.6 | 9.2 | 74.4 |
| | | WTH | Control | 32.5 | 2.2 | 23.0 | 41.9 |
| | | | Ash | 27.9 | 1.7 | 20.7 | 35.1 |
| | | | P-fertiliser | 36.1 | 6.1 | 10.0 | 62.3 |

| Hypothesis | Variable | Biomass Harvesting (BH) | Compensation Methods (CM) | Mean value | Standard Error | Lower confidence limit | Upper confidence limit |
|---|---|---|---|---|---|---|---|
| H4 *Soil acidity* | pH-H2O (unitless) | SOH | Control | 4.2 | 0.11 | 3.8 | 4.7 |
| | | | Ash | 4.6 | 0.11 | 4.2 | 5.1 |
| | | | P-fertiliser | 4.4 | 0.11 | 4.0 | 4.9 |
| | | AAH | Control | 4.8 | 0.11 | 4.3 | 5.2 |
| | | | Ash | 4.8 | 0.11 | 4.4 | 5.3 |
| | | | P-fertiliser | 4.5 | 0.11 | 4.0 | 4.9 |
| | | BAH | Control | 4.2 | 0.13 | 3.6 | 4.8 |
| | | | Ash | 4.5 | 0.11 | 4.1 | 5.0 |
| | | | P-fertiliser | 4.8 | 0.11 | 4.3 | 5.2 |
| | | WTH | Control | 4.4 | 0.11 | 4.0 | 4.9 |
| | | | Ash | 4.6 | 0.11 | 4.2 | 5.1 |
| | | | P-fertiliser | 4.5 | 0.11 | 4.0 | 4.9 |

| Hypothesis | Variable | Biomass Harvesting (BH) | Compensation Methods (CM) | Mean value | Standard Error | Lower confidence limit | Upper confidence limit |
|---|---|---|---|---|---|---|---|
| H4 *Soil acidity* | CEC base saturation (unitless) | SOH | Control | 0.40 | 0.04 | 0.25 | 0.56 |
| | | | Ash | 0.51 | 0.04 | 0.35 | 0.67 |
| | | | P-fertiliser | 0.38 | 0.04 | 0.22 | 0.54 |
| | | AAH | Control | 0.34 | 0.04 | 0.18 | 0.50 |
| | | | Ash | 0.43 | 0.04 | 0.27 | 0.59 |
| | | | P-fertiliser | 0.39 | 0.04 | 0.23 | 0.55 |
| | | BAH | Control | 0.42 | 0.04 | 0.26 | 0.58 |
| | | | Ash | 0.51 | 0.04 | 0.35 | 0.67 |
| | | | P-fertiliser | 0.35 | 0.04 | 0.19 | 0.51 |
| | | WTH | Control | 0.39 | 0.04 | 0.23 | 0.55 |
| | | | Ash | 0.40 | 0.04 | 0.24 | 0.56 |
| | | | P-fertiliser | 0.39 | 0.04 | 0.23 | 0.55 |

| Hypothesis | Variable | Biomass Harvesting (BH) | Compensation Methods (CM) | Mean value | Standard Error | Lower confidence limit | Upper confidence limit |
|---|---|---|---|---|---|---|---|
| H5 | soil Mn | SOH | Control | 1.18 | 0.15 | 0.52 | 1.80 |
| *Trace* | (mg/kg) | | Ash | 2.11 | 0.35 | 0.62 | 3.60 |
| *metals* | NH4-NO3 | | P-fertiliser | 1.48 | 0.30 | 0.21 | 2.70 |
| | extraction | AAH | Control | 1.12 | 0.08 | 0.76 | 1.50 |
| | | | Ash | 1.49 | 0.24 | 0.45 | 2.50 |
| | | | P-fertiliser | 1.16 | 0.12 | 0.65 | 1.70 |
| | | BAH | Control | 1.38 | 0.33 | -0.04 | 2.80 |
| | | | Ash | 1.69 | 0.05 | 1.46 | 1.90 |
| | | | P-fertiliser | 1.00 | 0.06 | 0.73 | 1.30 |
| | | WTH | Control | 1.00 | 0.05 | 0.77 | 1.20 |
| | | | Ash | 0.95 | 0.22 | 0.02 | 1.90 |
| | | | P-fertiliser | 1.04 | 0.17 | 0.33 | 1.80 |

| Hypothesis | Variable | Biomass Harvesting (BH) | Compensation Methods (CM) | Mean value | Standard Error | Lower confidence limit | Upper confidence limit |
|---|---|---|---|---|---|---|---|
| H5 | soil Zn | SOH | Control | 0.53 | 0.03 | 0.39 | 0.66 |
| *Trace* | (mg/kg) | | Ash | 0.68 | 0.13 | 0.12 | 1.24 |
| *metals* | NH4-NO3 | | P-fertiliser | 0.58 | 0.05 | 0.36 | 0.80 |
| | extraction | AAH | Control | 0.33 | 0.03 | 0.19 | 0.46 |
| | | | Ash | 0.60 | 0.11 | 0.15 | 1.06 |
| | | | P-fertiliser | 0.50 | 0.05 | 0.28 | 0.72 |
| | | BAH | Control | 0.46 | 0.03 | 0.33 | 0.60 |
| | | | Ash | 0.59 | 0.11 | 0.13 | 1.04 |
| | | | P-fertiliser | 0.36 | 0.05 | 0.14 | 0.58 |
| | | WTH | Control | 0.42 | 0.03 | 0.29 | 0.56 |
| | | | Ash | 0.46 | 0.11 | 0.01 | 0.92 |
| | | | P-fertiliser | 0.47 | 0.05 | 0.25 | 0.69 |

| Hypothesis | Variable | Biomass Harvesting (BH) | Compensation Methods (CM) | Mean value | Standard Error | Lower confidence limit | Upper confidence limit |
|---|---|---|---|---|---|---|---|
| H5 | soil Cd | SOH | Control | 8.5 | 1.1 | 4.0 | 13.1 |
| *Trace* | (µg/kg) | | Ash | 10.2 | 1.4 | 4.2 | 16.3 |
| *metals* | NH4-NO3 | | P-fertiliser | 9.1 | 0.6 | 6.4 | 11.7 |
| | extraction | AAH | Control | 7.9 | 1.1 | 3.4 | 12.5 |
| | | | Ash | 9.0 | 1.4 | 2.9 | 15.1 |
| | | | P-fertiliser | 8.6 | 0.6 | 6.0 | 11.3 |
| | | BAH | Control | 8.5 | 1.1 | 4.0 | 13.1 |
| | | | Ash | 10.5 | 1.4 | 4.4 | 16.6 |
| | | | P-fertiliser | 7.8 | 0.6 | 5.1 | 10.4 |
| | | WTH | Control | 7.8 | 1.1 | 3.3 | 12.4 |
| | | | Ash | 7.7 | 1.4 | 1.6 | 13.8 |
| | | | P-fertiliser | 7.1 | 0.6 | 4.5 | 9.8 |
