## Supplementary Figures and Supplementary Tables for "Response of soil and vegetation in a warm-temperate Pine forest to intensive biomass harvests, phosphorus fertilisation, and wood ash application"

**Figure S1** – Theoretical representation of the spatial distribution of treatments within one experimental block (Split-plot design)

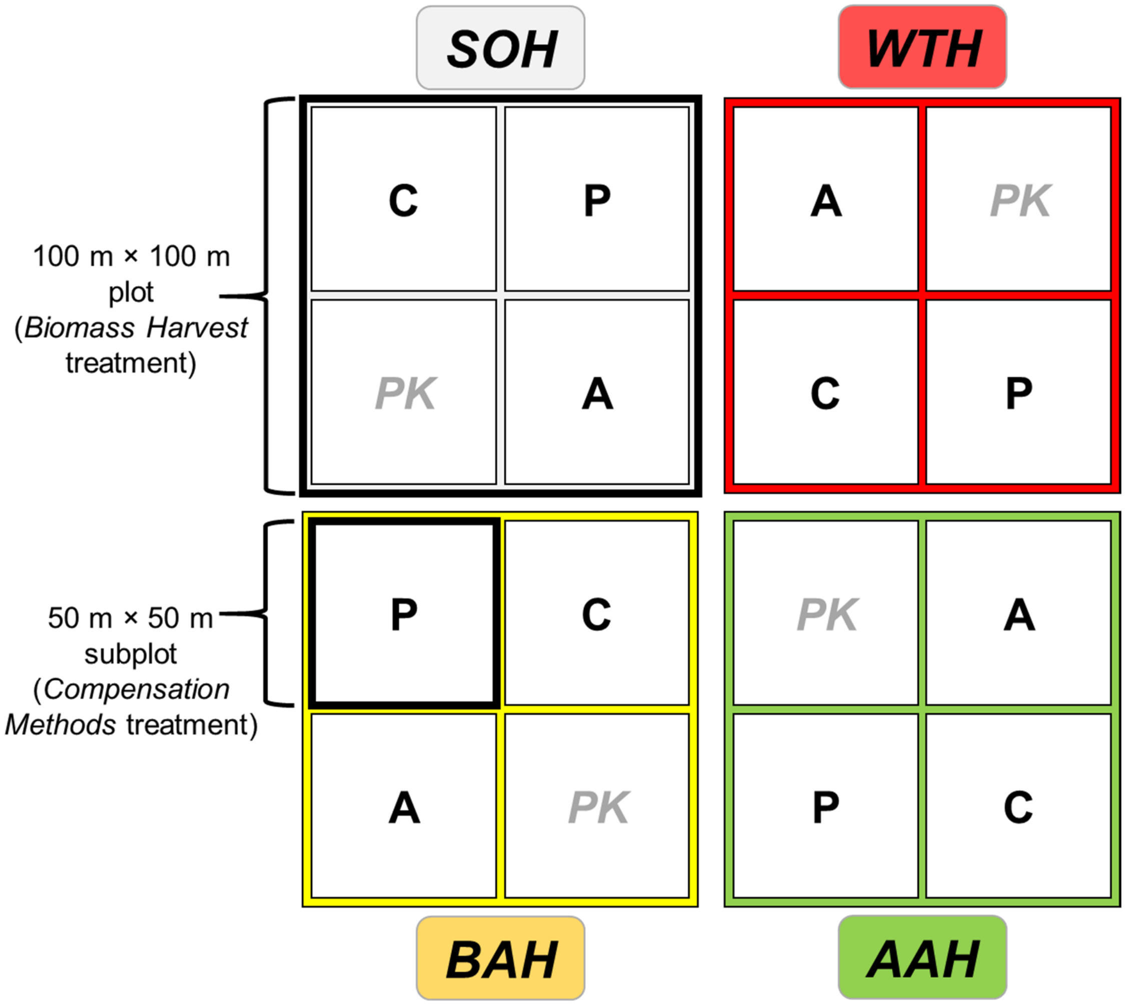


The experiment design was built using 3 complete blocks (example of 1 block in the scheme above). Each block was composed of 4 large plots, each dedicated to a different treatment of the *Biomass Harvest* factor. In turn, each plot was composed of 4 subplots, each dedicated to a different treatment of the *Compensation Methods* factor. The experiment was consequently based on 12 plots and 48 subplots. Because the PK treatment was not considered in the present study, only 36 subplots were sampled and monitored.

*Biomass Harvesting* treatments:

1- Stem-Only Harvest (SOH = stems; conventional harvest),
2- Aboveground Additional Harvest (AAH = stems + branches + foliage),
3- Belowground Additional Harvest (BAH = stems + stumps + roots),
4- Whole-Tree Harvest (WTH = stems + branches + foliage + stumps + roots).

*Compensation Methods* treatments:

1- Control: no nutrient application (C),
2- Phosphorus fertilisation (P),
*3- Phosphate-potassium fertilisation (PK; not considered in the present study),*
4- Wood Ash application (A).

**Figure S2** – Time sequence of the study

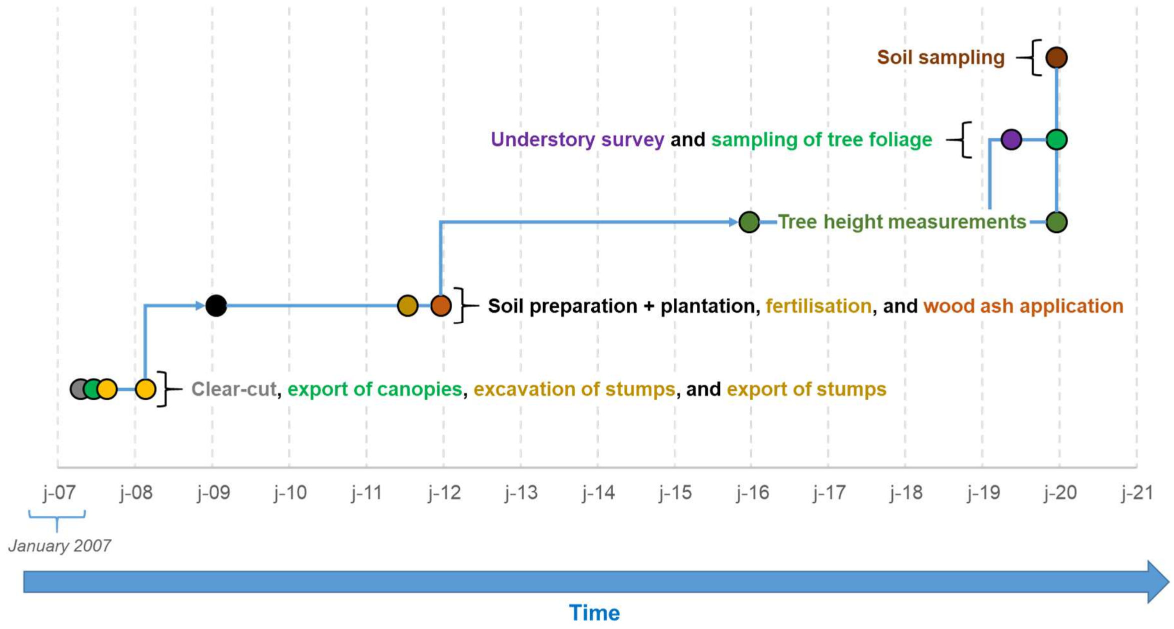

**Figure S3** – Different stages during the harvest of tree biomass

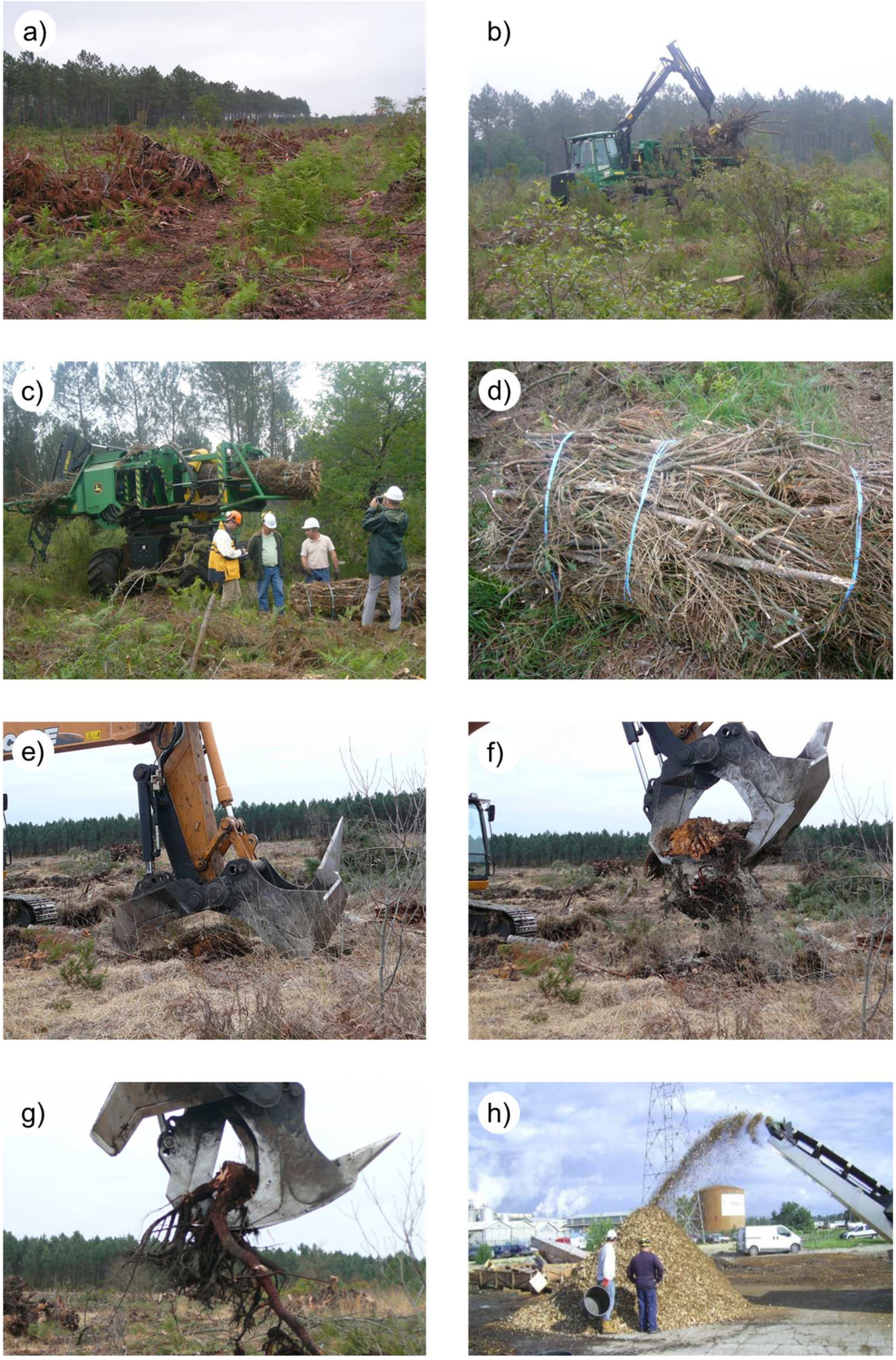


a) Tree canopies after two months of *in situ* drying; b) Harvest of branches with their remaining twigs and needles; c) Compaction of branches before the formation of a bundle; d) Zoom of a bundle, consisting of branches, twigs and needles; e) Machine used to extract stumps and roots. The "*horn*" is used to break the major roots directly in the soil and before extraction (not shown); f) The "*jaws*" are used to extract the stump from the soil, and g) to fragmentise stumps; h) chipping biomass samples (stump+roots biomass).

**Figure S4** – Field campaign for measuring understorey biomass based on phytovolume surveys

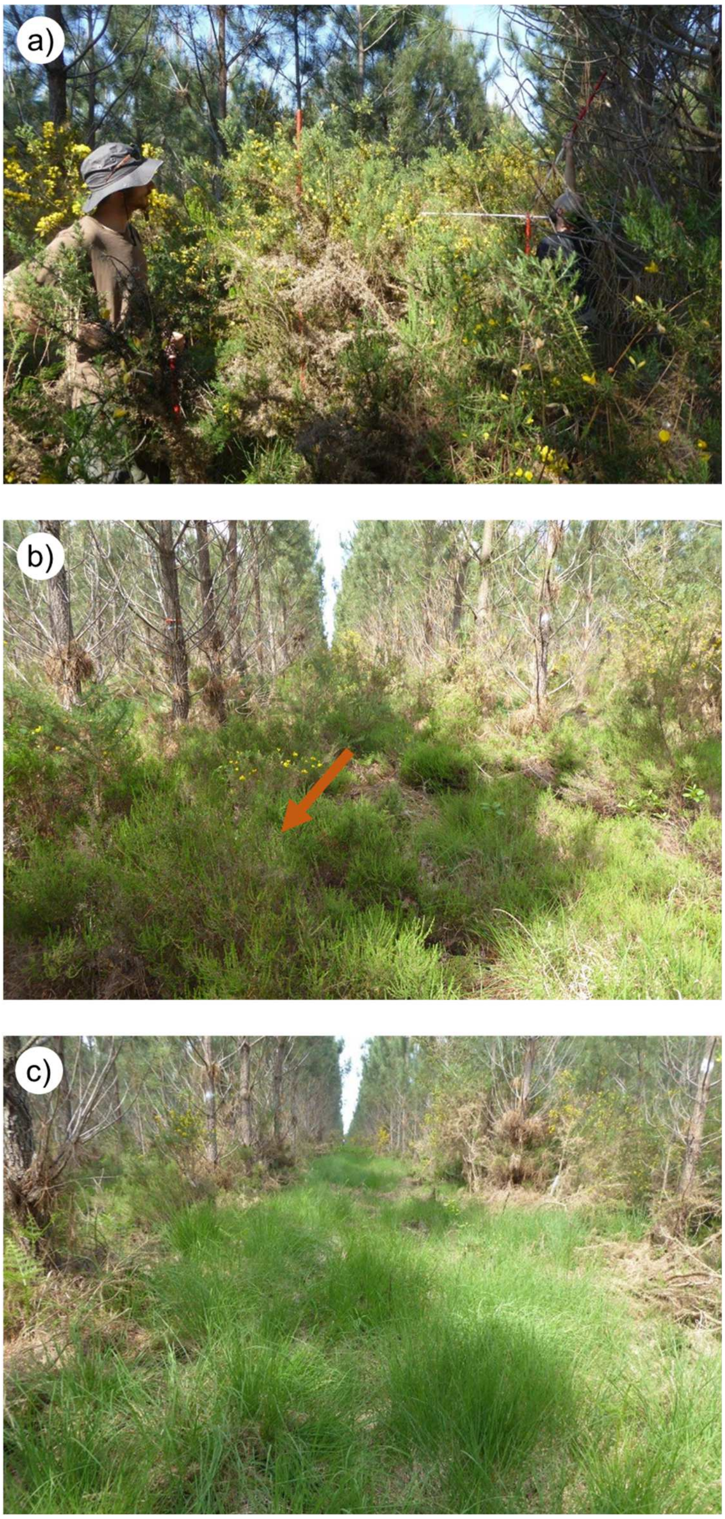


a) Two operators during the *phytovolume* estimates in a measuring area dominated by common gorse (*Ulex europaeus*), a local spiny shrub which is a N-fixer species; b) Area (indicated by an arrow) with predominant presence of the ericaceous species *Calluna vulgaris*; c) Area dominated by *Molinia caerulea*, a local herbaceous species which is a deciduous perennial. The operators are also authors of this article and consent the publication of these photographs.

**Figure S5** – Nutrient input-output budgets of major nutrients (N, P, K, Ca, Mg), for all treatments, and considering three dates after plantation of trees

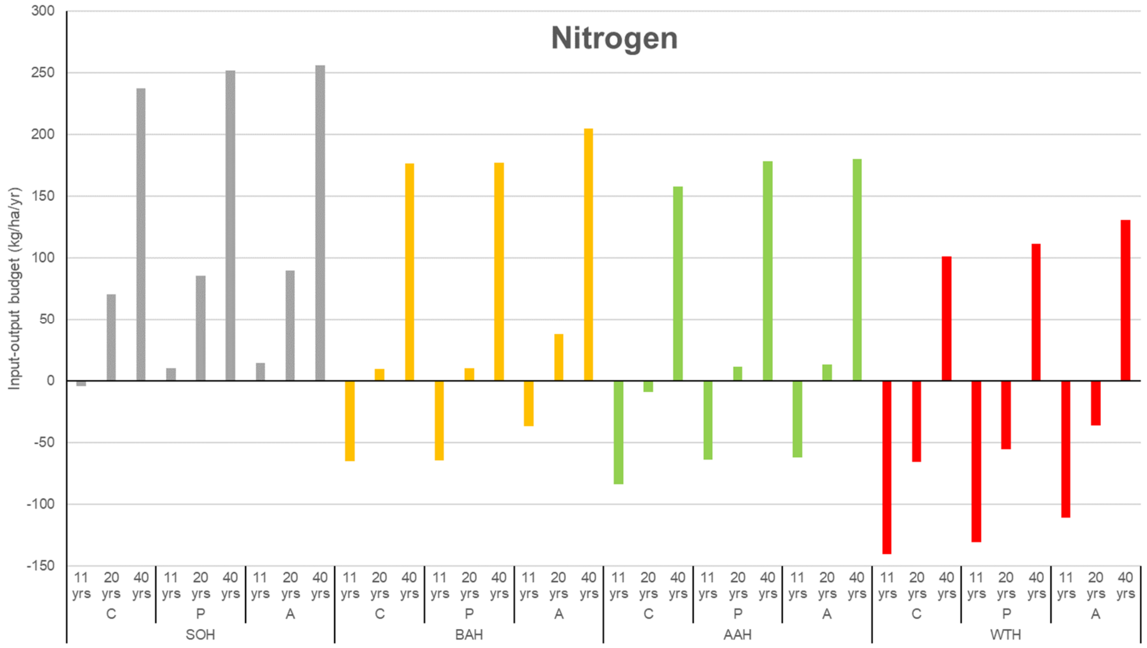


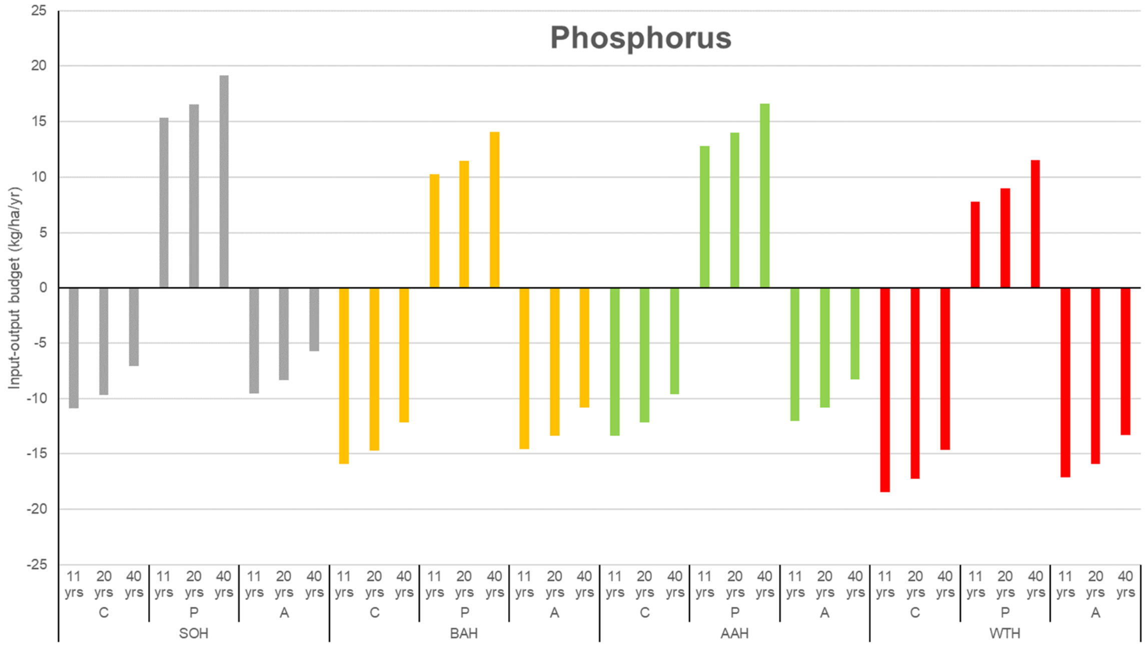

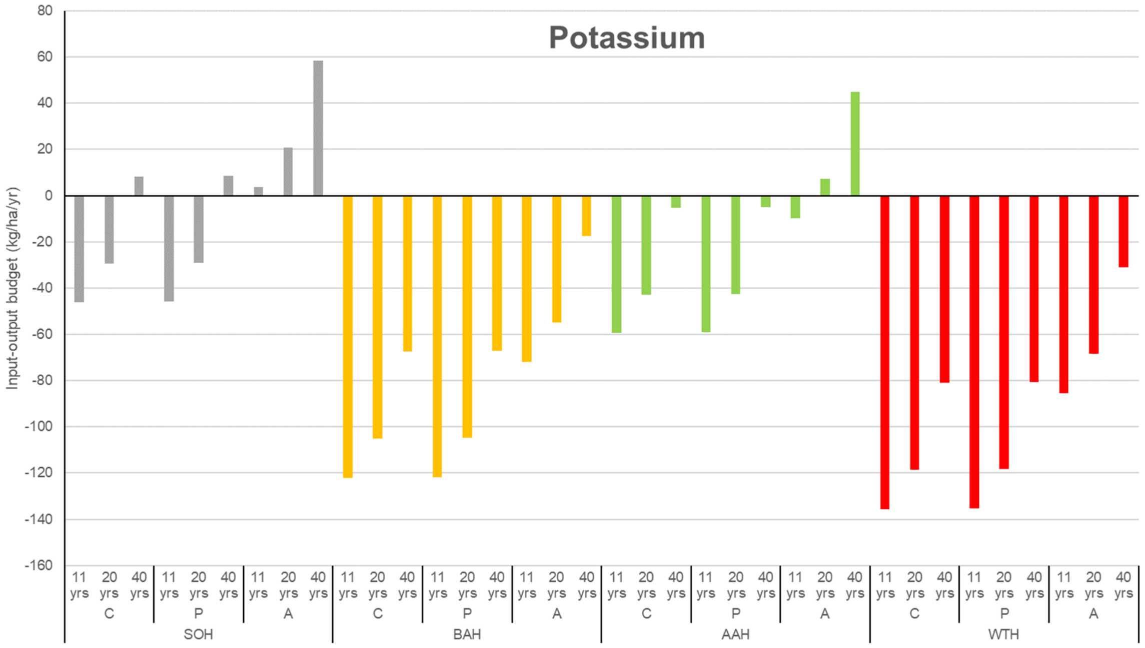
Potassium
Input-output budget (kg/ha/yr)
80
60
40
20
0
-20
-40
-60
-80
-100
-120
-140
-160
11 yrs
20 yrs
40 yrs
C
P
A
SOH
BAH
AAH
WTH

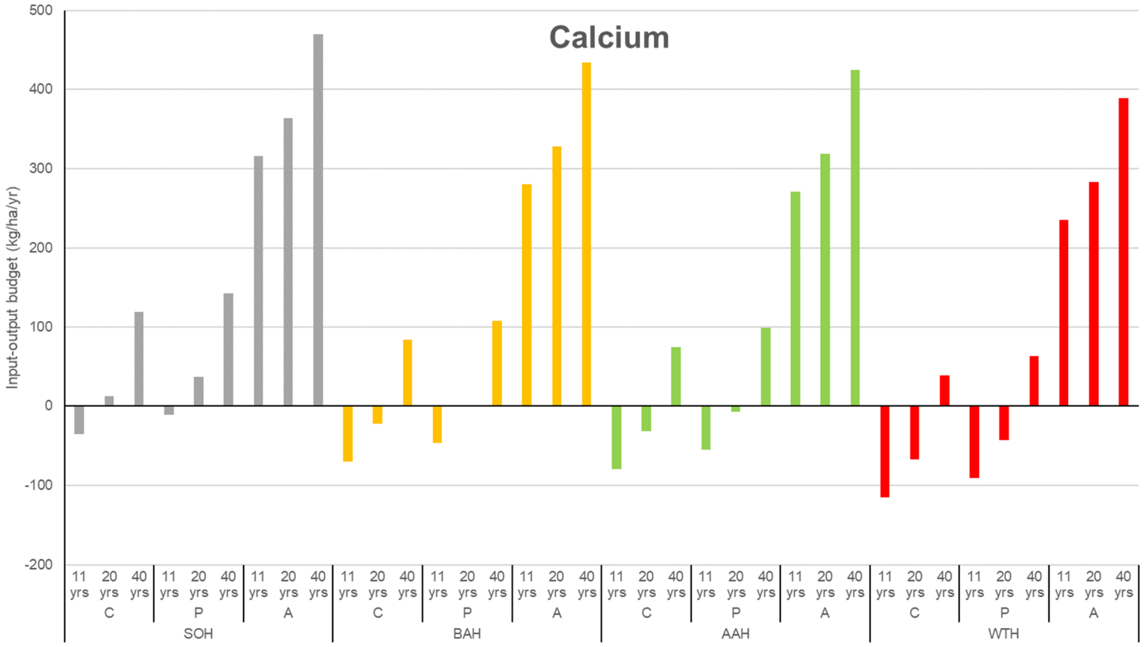
Calcium
Input-output budget (kg/ha/yr)
500
400
300
200
100
0
-100
-200
11 yrs
20 yrs
40 yrs
C
P
A
SOH
BAH
AAH
WTH

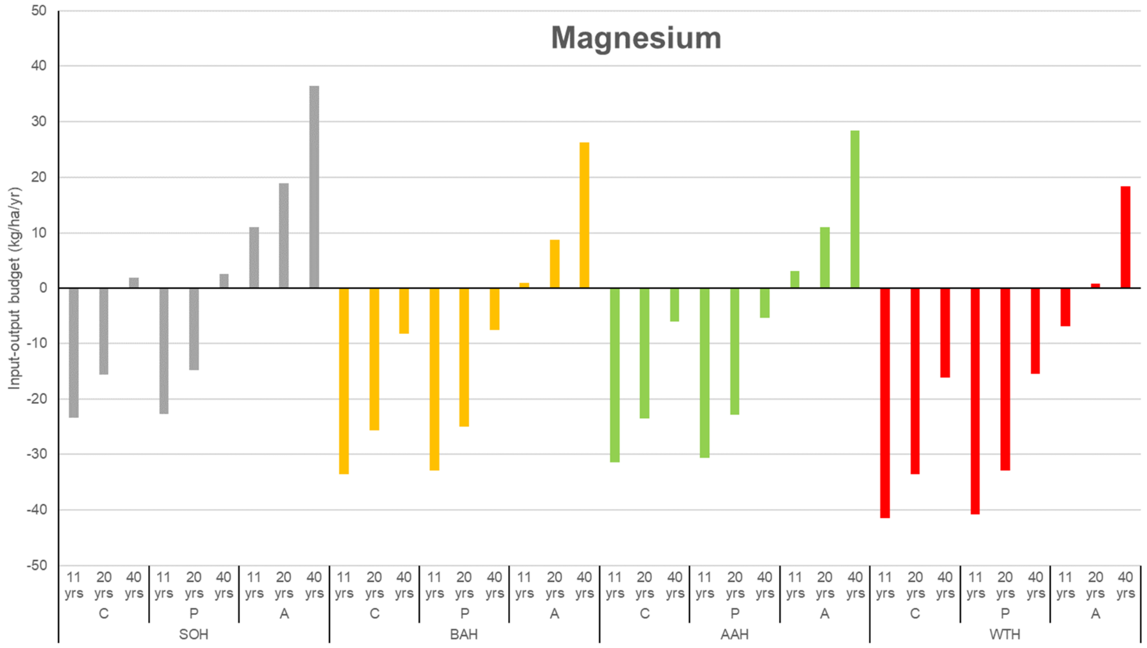


The input-output budgets take into account: atmospheric deposition, nitrogen symbiotic, fixation, weathering of soil minerals, fertilisation and wood ash application, biomass export, and deep seepage (see Methods).

Treatments considered were: stem-only harvest (SOH), aboveground additional harvest (AAH), belowground additional harvest (BAH), whole-tree harvest (WTH), combined with phosphorus fertilisation (P), wood ash application (A), or a control without any input (C).

Budgets are calculated for forest stands of 11 years-old (corresponding to the age of trees at most sampling dates), and for durations of the management rotation of 20 years or 40 years (corresponding to intensive forestry, or conventional forestry).

To enable comparisons with soil stocks ($kg_{-nutrient}$ $ha^{-1}$), nutrient pools are also presented for the 0-15 cm soil layer:

Total content values: N = 2051, P = 157, K = 7101, Ca = 1278, Mg = 268 kg $ha^{-1}$.

Available forms: P = 40, K = 32, Ca = 232, Mg = 52 kg $ha^{-1}$.

Values for the whole soil profile (0-120 cm), and for the same kind of forest ecosystems, are available in Augusto *et al.* (2010; Table 5).

**Table S1** – Composition of the wood ash applied during the field experiment

| Elements | Unit | Local wood ash used | Literature range |
|---|---|---|---|
| N | | < 10 | < 1 |
| P | | 0.27 | < 15 |
| K | Nutrients (kg $Mg^{-1}$) | nd | 2-130 |
| Ca | | 70.1 | 40-350 |
| Mg | | 6.9 | 3-25 |
| B | | 17 | 10-300 |
| Cu | | 37 | 15-300 |
| Fe | Micronutrients (g $Mg^{-1}$) | 5,340 | 5,000-20,000 |
| Mn | | 248 | < 30,000 |
| Mo | | < 4 | < 1-100 |
| Zn | | 148 | 15-2,200 |
| Al | | 11,500 | 10,000-30,000 |
| Cd | | 1.0 | < 1-25 |
| Cr | Non-essential elements (g $Mg^{-1}$) | 17 | 10-250 |
| Ni | | 10 | 6-200 |
| Pb | | 15 | 15-650 |

The ranges of values found in the literature are from a meta-analysis (Augusto et al. 2008). The quantity of nutrients applied in the field could be estimated based on the dose of wood ash in the experimental design (5 Mg $ha^{-1}$): for instance, 351 and 35 kg $ha^{-1}$ of Ca and Mg.

**Table S2** – Biomass and nutrient content of the stand before the harvests (**a**), and in the exports (**b**)

**a**) Standing biomass and its nutrient content ⇔ *available biomass and nutrients before harvesting operations*

| *Tree compartments* | Biomass (Mg ha$^{-1}$) | N (kg ha$^{-1}$) | P (kg ha$^{-1}$) | K (kg ha$^{-1}$) | Ca (kg ha$^{-1}$) | Mg (kg ha$^{-1}$) |
|---|---|---|---|---|---|---|
| Foliage | 5.28 ± 0.10 | 55.0 ± 1.0 | 2.21 ± 0.04 | 11.3 ± 0.2 | 21.0 ± 0.4 | 6.19 ± 0.11 |
| Twigs | 4.46 ± 0.13 | 29.0 ± 0.8 | 1.28 ± 0.04 | 7.2 ± 0.2 | 31.0 ± 0.9 | 3.60 ± 0.10 |
| Branches | 11.38 ± 0.25 | 49.0 ± 1.1 | 1.82 ± 0.04 | 9.8 ± 0.2 | 36.0 ± 0.8 | 6.40 ± 0.14 |
| Stembark | 12.99 ± 0.23 | 28.3 ± 0.5 | 1.69 ± 0.03 | 11.7 ± 0.2 | 31.7 ± 0.6 | 5.72 ± 0.10 |
| Stemwood | 104.93 ± 2.21 | 63.4 ± 1.3 | 10.49 ± 0.22 | 52.5 ± 1.1 | 52.5 ± 1.1 | 23.08 ± 0.49 |
| Stump and roots | 40.09 ± 0.72 | 85.9 ± 1.5 | 7.16 ± 0.13 | 107.4 ± 1.9 | 50.1 ± 0.9 | 14.32 ± 0.26 |
| Total | 179.14 ± 3.53 | 310.6 ± 5.8 | 24.66 ± 0.47 | 199.8 ± 3.7 | 222.2 ± 4.3 | 59.30± 1.14 |

**b**) Realised biomass harvests and their concomitant nutrient exports (mean values) ⇔ *biomass and nutrients that were actually exported*

| *Biomass Harvests* | Biomass (Mg ha$^{-1}$) | N (kg ha$^{-1}$) | P (kg ha$^{-1}$) | K (kg ha$^{-1}$) | Ca (kg ha$^{-1}$) | Mg (kg ha$^{-1}$) |
|---|---|---|---|---|---|---|
| SOH | 115.32 | 86.0 | 11.84 | 61.9 | 77.9 | 27.66 |
| AAH | 126.49 | 149.9 | 14.36 | 75.3 | 122.4 | 35.57 |
| BAH | 143.66 | 146.8 | 16.90 | 137.8 | 113.3 | 37.78 |
| WTH | 154.83 | 210.6 | 19.42 | 151.3 | 157.9 | 45.69 |

SOH = stem-only harvest; AAH = aboveground additional harvest; BAH = belowground additional harvest; WTH = whole-tree harvest.
The values of realised harvests (Table S2b) for biomass and N-P-K are illustrated in Figure 1 in the main text.

**Table S3** – Tree size (CBH, VOL, Height) and growth (*i*H-4yrs) as affected by biomass harvest and compensation methods

| | CBH (cm) | $VOL_{stem}$ ($dm^3$ $tree^{-1}$) | Height (m) | *i*H-4yrs (m) |
|---|---|---|---|---|
| *Biomass Harvesting* | | | | |
| SOH | 42.1±0.6 | 57±2 | 8.57±0.13 | 3.80±0.07 |
| AAH | 40.9±0.6 | 51±2 | 8.23±0.13 | 3.87±0.07 |
| BAH | 41.1±0.6 | 53±2 | 8.38±0.13 | 3.82±0.07 |
| WTH | 40.1±0.6 | 50±2 | 8.39±0.13 | 3.77±0.07 |
| *Compensation Methods* | | | | |
| Control | 41.2±0.5 | 53±2 | 8.41±0.10 | 3.78±0.06 |
| Phosphorus | 39.6±0.5 * | 47±2 * | 8.14±0.10 * | 3.73±0.06 |
| Ash | 42.4±0.5 | 57±2 (*) | 8.62±0.10 | 3.93±0.06 (*) |

SOH = stem-only harvest; AAH = aboveground additional harvest; BAH = belowground additional harvest; WTH = whole-tree harvest. Control = no nutrient application. CBH = stem circumference at breast height (≈ 1.3 m); $VOL_{stem}$ = stem volume; *i*H-4yrs = height growth during the last four years. Trees were measured at 11 years old. Treatments with an asterisk differ at $P<0.05$ (for $P$ value < 0.10, the symbol is in brackets) from their reference treatment (SOH or Control, respectively; see Methods). Mean values of the Biomass Harvesting treatments (n = 3 blocks) were calculated by pooling the Compensation Methods subplots. Mean values of the Compensation Methods treatments were calculated based on 12 replicates (3 blocks × 4 BH treatments).

**Table S4** – Soil composition in water soluble nutrients and micronutrients as affected by biomass harvest and compensation methods

| | P ($\mu$g g$^{-1}$) | K ($\mu$g g$^{-1}$) | Ca ($\mu$g g$^{-1}$) | Mg ($\mu$g g$^{-1}$) | Fe ($\mu$g g$^{-1}$) | Mn ($\mu$g kg$^{-1}$) | Zn ($\mu$g kg$^{-1}$) |
|---|---|---|---|---|---|---|---|
| *Biomass Harvesting* | | | | | | | |
| SOH | 1.4±0.2 | 7.3±0.3 | 2.6±0.5 | 1.9±0.2 | 2.0±0.3 | 32±5 | 9.4±1.3 |
| AAH | 0.6±0.1 * | 6.2±0.3 | 2.5±0.2 | 1.9±0.2 | 2.6±0.3 | 40±4 | 7.6±1.0 |
| BAH | 1.4±0.1 | 5.7±0.3 * | 2.7±0.3 | 1.5±0.2 | 1.9±0.3 | 32±3 | 6.9±0.6 |
| WTH | 1.3±0.3 | 5.7±0.3 * | 2.1±0.2 | 1.5±0.2 | 1.6±0.3 | 20±3 | 6.5±0.7 |
| *Compensation Methods* | | | | | | | |
| Control | 1.2±0.2 | 5.8±0.3 | 2.6±0.2 | 1.6±0.1 | 1.9±0.4 | 27±3 | 6.8±0.6 |
| Phosphorus | 1.2±0.2 | 6.4±0.3 | 2.1±0.4 * | 1.7±0.1 | 2.1±0.3 | 28±2 | 7.6±0.6 |
| Ash | 1.1±0.1 | 6.5±0.3 (*) | 2.6±0.2 | 1.8±0.1 | 2.1±0.2 | 39±4 * | 8.8±0.6 * |

SOH = stem-only harvest; AAH = aboveground additional harvest; BAH = belowground additional harvest; WTH = whole-tree harvest. Control = no nutrient application. Treatments with an asterisk differ at $P$<0.05 (for $P$ value < 0.10, the symbol is in brackets) from their reference treatment (SOH or Control, respectively; see Methods). Mean values of the Biomass Harvesting treatments (n = 3 blocks) were calculated by pooling the Compensation Methods subplots. Mean values of the Compensation Methods treatments were calculated based on 12 replicates (3 blocks × 4 BH treatments).
It should be noted that the units differ among the elements.

**Table S5** – Initial expected consequences of intensive biomass harvesting and wood ash application on the studied ecosystem (temperate oligotrophic forest)

(➔: means "has no effect on…"; ➘: means "decreases..."; ➚: means "increases..."; ❌: means "hypothesis was not validated"; ✅: means "hypothesis was validated"; ±: means "hypothesis received mixed support")

| Ecosystem trait | | Intensive harvests | Wood ash |
|---|---|---|---|
| Soil fertility (phosphorus) {hypothesis H1} | *Expected* | ➘ (aggravated P-limitation) | ➔ (or ➚) soil P |
| | *Observed* | ✅ | ✅ (➔) |
| Tree growth (carbon sequestration) {H2} | *Expected* | ➘ tree growth | ➔ (or ➚) tree growth |
| | *Observed* | ± (➘ trend only) | ✅ (➚ trend) |
| Soil Organic Matter (C and N) {H3} | *Expected* | ➘ SOM | ➔ SOM |
| | *Observed* | ✅ | ❌ (➘) |
| Soil acidity (pH, base saturation) {H4} | *Expected* | ➘ acido-basic status | ➚ acido-basic status |
| | *Observed* | ❌ (➔) | ✅ |
| Trace metals in ecosystem {H5} | *Expected* | ➔ trace metals | ➔ or ➘ bioavailability at moderate ash dose |
| | *Observed* | ✅ | ✅ (➔) |
